# First crystal structure of a non-canonical amino acid linked to a paramagnetic lanthanide tag facilitates protein structure determination using NMR-derived restraints

**DOI:** 10.1101/2022.04.10.487812

**Authors:** Elleansar Okwei, Soumya Ganguly, Heather L. Darling, Joel M. Harp, Alican Gulsevin, Irene Coin, Hassane Mchaourab, Kaitlyn Ledwitch, Georg Kuenze, Jens Meiler

## Abstract

Site-directed spin labeling of proteins via non-canonical amino acids (ncAAs) is a non-traditional method for the measurement of pseudocontact shifts (PCSs) by nuclear magnetic resonance (NMR) spectroscopy. PCSs provide long-range distance and orientational information between a paramagnetic center and protein nuclei that can be used as restraints for computational structural modeling techniques. Here, we present the first experimental structure of an ncAA chemically linked to a lanthanide tag conjugated to the protein, T4-Lysozyme (T4L). T4L was crystallized with a cyclen-based C3 tag coordinated to the paramagnetic ion terbium (Tb^3+^). The paramagnetic C3-lanthanide tag generated PCSs measured at four different ncAA sites. We show that the addition of these restraints improves structure prediction protocols for T4L using the RosettaNMR framework. Generated models provide insight into T4L conformational flexibility sampled in solution. This integrative modeling protocol is readily transferable to larger proteins. Methods to predict protein structures are advancing into an exciting arena such that reliable experimental data will play important roles for evaluating the biophysical relevance of predicted structural models. Our contribution here caters to the growing interest in using ncAAs for a range of biophysical studies, and these methods can be readily transferred to larger protein systems of interest.

## Introduction

Paramagnetism in NMR was observed as early as the 1960s when Shulman and coworkers noticed in a ^1^H NMR spectrum of paramagnetic heme proteins that the ^1^H signals of the heme groups were shifted relative to the signals of the rest of the protein (Shulman et al., 1969). Several studies since have taken advantage of the paramagnetic center in metalloproteins (Lee and Sykes, 1980a) or substituting diamagnetic with paramagnetic metal ions such as lanthanides (Griffin et al., 1975; Lee and Sykes, 1980b), to intentionally generate paramagnetic effects in NMR. Lanthanides are suitable for paramagnetic NMR because of their versatility and the pronounced effects they produce in NMR spectra. Since most proteins do not possess a paramagnetic center, studying proteins via paramagnetic NMR spectroscopy has been hindered by 1) challenges associated with introducing a paramagnetic moiety into a protein and 2) availability of suitable paramagnetic probes to generate accurate and reliable paramagnetic NMR restraints.

Typically, a paramagnetic probe is attached to the protein of interest through a cysteine residue via a disulfide or thioether linkage, which affects NMR signals of nearby nuclei in a distance-dependent manner (Battiste and Wagner, 2000; Ikegami et al., 2004; Peters et al., 2011). However, important classes of proteins may have disulfide bridges that play a role in protein stability or function and thus, approaches that require mutation of cysteines are not always possible. An alternative approach is the introduction of a non-canonical amino acid (ncAA), which does not require the removal of all native cysteines and re-introduction of single cysteine mutants for labeling. Genetic code expansion allows ncAA incorporation into proteins for biophysical studies (Young et al., 2010). An ncAA can be introduced site-specifically (Wals and Ovaa, 2014), and usually contains a functional group orthogonal to native protein chemistry. Para-azido-phenylalanine (pAzF) is an ncAA that has an azide group that can be reacted with an alkyne via a copper(I)-catalyzed azide-alkyne cycloaddition (CuAAC) reaction (Hong et al., 2009). The Otting group demonstrated this with a cyclen-based label, a nitrilotriacetic acid-based label, and an iminodiacetic acid-based label that was developed for site-specific labeling to obtain paramagnetic NMR restraints (Loh et al., 2015; Loh et al., 2013). In this work, we present the first determined crystal structure of a protein site-specifically conjugated via an ncAA to the cyclen-based C3 lanthanide label.

NMR spectroscopy has contributed to exceptional advances in investigating protein structure and dynamics (Arora and Tamm, 2001; Griesinger et al., 2004; Kuenze, 2019; Liang et al., 2006; MacKenzie et al., 1997; Meiler et al., 2000a; Meiler et al., 2000b, 2003; Oxenoid and Chou, 2005; Peti et al., 2002; Van Horn et al., 2009; Weiner et al., 2014). Paramagnetic restraints measured by NMR can complement or replace nuclear Overhauser effects (NOEs) for structure prediction. In addition to pseudocontact shifts (PCSs) discussed in this work, other effects such as paramagnetic relaxation enhancements (PREs), and residual dipolar couplings (RDCs) (Bax, 2003; Howell et al., 2005; Opella and Marassi, 2004; Sanders and Sonnichsen, 2006; Tamm, 2006; Tjandra and Bax, 1997; Tolman et al., 1995) can be observed when a paramagnetic center is present in the protein; although not all paramagnetic centers elicit all three types of restraints.

Attaching paramagnetic tags at multiple locations determines the positions of secondary structural elements within the protein fold by triangulation similar to the global positioning system (GPS) (Chen et al., 2011). This concept can be incorporated into computational frameworks for guiding structure prediction using experimental information (Kuenze, 2019). This is particularly beneficial in the case of measuring PCSs, which provides both distance and angular restraints for the protein of interest.

PCSs are one of the most useful sources of paramagnetic data. They depend on the polar coordinates r, θ, and φ of the nuclear spin with respect to the magnetic susceptibility Δχ tensor of the metal ion and the axial and rhombic components of the Δχ tensor (Bertini et al., 2002):

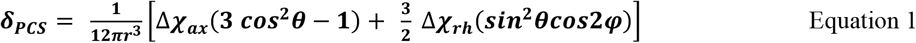

PCSs can be measured in two-dimensional heteronuclear single quantum coherence (HSQC) spectra of ^15^N-labeled proteins as altered chemical shifts. Depicting the PCSs as isosurfaces onto the 3D structure of the protein gives an alternative representation of the Δχ tensor and the orientations of its principal axes. Thus, PCSs give access to a coordinate system that is tied to the protein molecule and centered at the metal ion. This reference frame allows positioning nuclear spins by their PCSs (Otting, 2010).

Sparse or incomplete experimental datasets integrated with computational modeling supplements missing experimental information with atomic detail. Stellar progress has been made in structure prediction of soluble proteins either *de novo* (starting from sequence) (Baek et al., 2021; Bradley et al., 2003; Bradley et al., 2005a; Bradley et al., 2005b; Jumper et al., 2021), or through comparative modeling that uses structural information from homologous proteins to the target protein (Song et al., 2013). These methods have also been applied to integral membrane proteins (Barth et al., 2007; Barth et al., 2009; Bender et al., 2020; Li et al., 2019; Li et al., 2016, 2017; Weiner et al., 2013; Yarov-Yarovoy et al., 2006), but the size and complexity of these proteins limit the success of these methods in the absence of experimental data. More recently, some modeling platforms performed well without experimental data (Baek et al., 2021; Jumper et al., 2021), however experimental evaluation is key to interpreting the biophysical relevance of predicted models. Thus, there is added benefit to including experimental data to guide computational methods for structure prediction. For example, LmrP was determined to high accuracy with Alphafold2 in one model, but an alternative model with a good score was in disagreement with the experimentally-derived structure. Experimental data from electron paramagnetic resonance (EPR) spectroscopy proved useful in validating that the predicted structural conformation was biologically-relevant (Hegedus et al., 2022).

Integrative structural modeling leverages information from multiple biophysical and biochemical sources to construct a holistic, atomic-detail model consistent with all available data (Russel et al., 2012; Xia et al., 2018). It led to the identification of structures of complex protein systems (Demers et al., 2014; Robinson et al., 2015). Integrative modeling connects structural and experimental data in a quantitative way to inform more about the biological system than possible with individual methods alone. Consequently, approaches that combine computational techniques with experimental techniques (e.g. X-ray crystallography, NMR, electron microscopy, EPR) can be used to address the shortcomings of individual experimental methods and are superior to computational models that are not validated by experimental data. Structural models of complex systems are increasingly characterized using integrative methods and are disseminated to the public through PDB-Dev, which serves as a prototype archiving system for integrative structures (Berman et al., 2019; Burley et al., 2017; Sali et al., 2015; Vallat et al., 2019; Vallat et al., 2018). Beyond predicting protein structures accurately, combining the power of integrative modeling with experimental NMR data (i.e. PCSs) is advantageous for investigating flexible, heterogeneous systems like protein-ligand and protein-nucleic acid systems, protein complexes, and protein conformational dynamics (Kuenze, 2019).

To effectively measure PCSs for a biological system, careful consideration must be given to the paramagnetic label of choice and the method of introduction to the protein system. Increased interest in lanthanide-induced paramagnetic effects has led to the development of a plethora of lanthanide-binding tags (LBTs) that can be incorporated into proteins. Many of the developed lanthanide tags are quite bulky and could potentially disturb the protein structure. It has remained elusive how the conformation of such tags changes when attached to a protein, and if these changes perturb protein structure. Furthermore, crystallographic evidence for the tag conformation can improve protein structure determination using NMR restraints by providing a precise location of the metal ion position. Here, we determined the first crystal structure of an ncAA attached to the C3-Tb^3+^ LBT, bound to a model protein system, T4L. This crystallographic model will help support PCS-guided protein integrative structural modeling.

Our group recently introduced a comprehensive framework into the Rosetta suite that uses NMR restraints derived from paramagnetic labeling (Kuenze, 2019). In this current work, we describe an experimental approach for the collection of paramagnetic data to complement the computational protocol. Specifically, we demonstrate that this strategy provides new structural and conformational insight to the flexible bacteriophage T4L using PCSs measured at multiple sites with the C3-lanthanide tag. Crystallographic evidence of label position and orientation conjugated to pAzF were used together with PCSs obtained from NMR for computational modeling of T4L. The results presented here demonstrate the utility of PCSs for guiding computational modeling of flexible proteins, and for analyzing conformational changes in solution.

T4L breaks down the peptidoglycan wall of the bacterial host of bacteriophage T4 and is a well-characterized protein that has served as a model system for various biophysical investigations. As seen in the mutant crystal structures available in the protein data bank (PDB), it contains two subdomains linked by an inter-helical domain [Figure 1A]. The amino-terminal domain comprises residues 15 to 59, the carboxy-terminal domain comprises residues 80 to 162, and residues 1 to 14 make up the hinge-bending region; the long-interhelical domain spans residues 60 to 79 (Zhang et al., 1995). Owing to its hinge-bending region, T4L has been observed to adopt multiple conformations (Faber and Matthews, 1990) - mainly a more compact closed state (Kuroki et al., 1993; Weaver and Matthews, 1987) or a more extended open state (Dixon et al., 1992) [Figure 1B]. Over 90% of the T4L crystal structures represented in the PDB are in the closed state, even though the hinge-bending motion suggests a wider range of conformations. Thus, other methods for analyzing the hinge-bending motion in solution or without crystal packing constraints have been employed. EPR spectra comparing free and substrate-bound T4L with nitroxide tags confirmed that the hinge-bending motion was associated with substrate binding and T4L catalysis in solution (Mchaourab et al., 1997). This is also supported through molecular dynamics simulations that suggest transitions between different conformations of T4L (Biggers et al., 2020; de Groot et al., 1998; Vallurupalli et al., 2016). Dipolar couplings measured in liquid crystalline media by solution NMR methods identified that the cleft between the domains is significantly larger in the average solution structure than what is observed in the X-ray structure of the ligand-free form of the protein (Goto et al., 2001). Through a site-specific lanthanide binding site, PCSs and RDCs were generated by attaching 4-mercaptomethyl dipicolinic acid (4MMDPA) via a disulfide bond to a single mutant of T4L. The data indicated that in solution and in the absence of substrate, the structure of T4L is on average more open than suggested by the closed conformation of the crystal structure of wild type T4L (Chen et al., 2016). Structure-based design calculations using relaxation-dispersion NMR and CS-Rosetta model building enabled studies of a solution structure of a minor and transiently formed state of a T4L mutant (Bouvignies et al., 2011).

**Figure 1.**
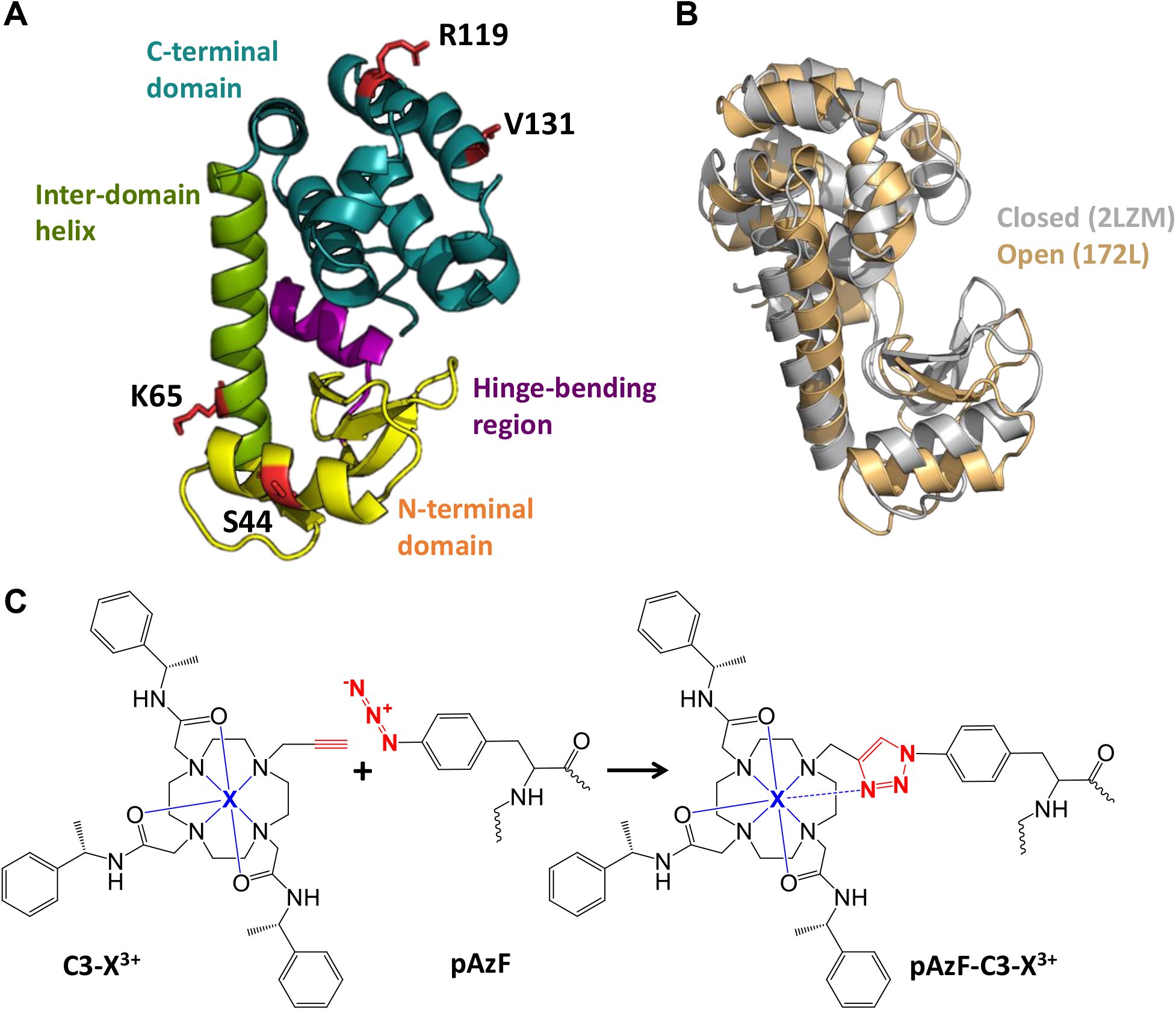
Mutants generated for site-directed lanthanide ion labeling (SDLL) in T4L for sparse PCS measurements. (A) T4L wild type (PDB ID: 2LZM) consists of an N-terminal domain (yellow) and C-terminal domain (teal) connected by a long α-helix (green). The motion in the hinge-bending region (purple) governs its flexibility and the ability of T4L to adopt multiple conformations. For broad coverage of the protein, four sites (shown as red sticks) were selected for labeling (S44, K65, R119, and V131). (B) The two pre-dominant conformations: closed (colored gray (PDB ID 2LZM) and open (yellow; PDB ID: 172L) are shown. (C) The cyclen-based lanthanide label (C3-X^3+^, where X^3+^ represents a lanthanide ion: Y^3+^, Yb^3+^, Tm^3+^, or Tb^3+^), was chosen for measuring sparse paramagnetic restraints. The reaction scheme (as seen in Loh et al, 2013) shows the alkyne moiety of the C3 label (red) reacting with the azide of p-azido-L-phenylalanine (pAzF), colored red, at the selected site in the protein using Cu(I)-catalyzed click chemistry. Each single pAzF mutant was produced and labeled for NMR studies. The chelated lanthanide is colored blue. Reaction scheme adapted with permission from *Bioconjugate Chem*. 2013, 24, 2, 260–268. Copyright 2013 American Chemical Society.”

In this work, we describe an experimental approach for the collection of paramagnetic data for use in the RosettaNMR computational framework developed in our group (Kuenze, 2019). We provide crystallographic evidence of ncAA-linked paramagnetic C3 tag and the position and orientation of the tag with respect to T4L. PCSs were collected at multiple tagging sites and used for experimentally-informed computational modeling of T4L in Rosetta. The utility of PCSs for guiding computational modeling of α-helical protein regions, full-length flexible proteins, and for studying protein conformations in solution is demonstrated. The experimental strategy described here resulted in a set of sparse restraints that improved full-length T4L structure prediction using the RosettaNMR framework. Additionally, the PCSs also encoded information related to the structural dynamics of T4L and allowed observation of a range of solution conformations not readily observed in the X-ray crystal structures available in the PDB.

## Results and Discussion

### Probing paramagnetic effects of a C3 lanthanide binding tag (LBT) conjugated to pAzF, at multiple sites using an alpha-helical protein as a model system

T4L was chosen as a model protein system for three reasons: 1) T4L structure is well-characterized (there are > 500 structures in the PDB) allowing straightforward comparison with computational and NMR results; 2) likelihood to crystallize in the presence of the bulky C3 tag, and 3) the helical domain of T4L (the inter-helical and C-terminal regions of T4L) would allow assessment of the ability of computational methods to predict structures of all-helical proteins and would have broader applications for systems like membrane proteins. An additional benefit of using T4L is that because it is flexible (e.g. in the N-terminal domain + hinge-bending region) and has been shown to adopt multiple conformations in solution, solution conformations encoded in the PCS data can be distinguished. Ultimately, using T4L as a model system facilitated an analysis of how well sparse PCS datasets guides computational modeling of not just the ordered α-helical regions, but also the full-length flexible protein. Our experimental approach provided a reliable PCS dataset that allowed re-determination of T4L structure and increased precision of computational modeling with a set of PCSs measured at multiple sites.

The T4L construct used here was the “pseudo-wildtype” (C54T/C97A) which has been shown to maintain the same structure and function as the wild type T4L (Matsumura and Matthews, 1989). Four sites were chosen (S44, K65, R119, or V131) for site-directed lanthanide spin labeling with three paramagnetic metals (i.e. Yb^3+^, Tm^3+^, Tb^3+^); the specific sites were solvent-exposed residues and have been employed in previous EPR experiments to study side chain dynamics (Mchaourab et al., 1999; Mchaourab et al., 1996). Furthermore, the selected sites are located on different α-helices spread over the protein sequence space such that the PCS datasets collected for these different sites are expected to carry complementary structural information.

T4L mutants were expressed with an amber stop codon (UAG) to replace residue Ser44, Lys56, Arg119 or Val131 (Figure 1A) with pAzF (referred to in the following as pAzF44, pAzF65, pAzF119, or pAzF131). Site-specific incorporation of pAzF was completed using the published pEVOL vector with the aminoacyl-tRNA synthetase for pAzF (pAzF-RS) (Young et al., 2010). With pAzF present in the protein, the C3-alkyne-lanthanide probe reacts with the azide group via a CuAAC click chemistry reaction (Loh et al., 2013) (Figure 1B).

### Structure determination of C3-lanthanide-labeled T4L

We crystallized and determined the structure for the pAzF65 mutant of T4L with C3-Tb^3+^ (T4L-pAzF65-C3-Tb^3+^) at a resolution of 1.89 Å. Crystal screening conditions and structure determination procedures are discussed in detail in the Methods section. The protein crystallized in space group P 43 21 2. Due to the presence of the anomalous scattering Tb^3+^ ion, anomalous diffraction data was analyzed. The programs SHELX C/D/E (Sheldrick, 2010) located one anomalous scattering atom, whose electron density is shown in Figure 2A. Single-wavelength anomalous dispersion (SAD) phasing was performed using the initial density map, then density was modified before building the final model. The C3 tag was modeled into its very prominent electron density (Figure 2B). For the final structure of T4L-pAzF65-C3-Tb^3+^ (Figure 2C), an R-work of 0.24 and R-free of 0.26 indicated a good measure of agreement between model and original diffraction data. A figure of merit (FOM) of 0.48 indicated superior phase quality for the single wavelength anomalous diffraction (SAD) data. Data collection and refinement statistics are provided in Table 1. The determined crystal structure has been deposited in the PDB under the accession code 7U4X.

**Table 1.** Data Collection and Refinement Statistics.

|  |  |
| --- | --- |
| Wavelength | 0.97857 Å |
| Resolution range | 38.82 - 1.786 (1.85 - 1.786)<sup>a</sup> |
| Space group | P 43 21 2 |
| Unit cell | 59.443 59.443 101.31 90 90 90 |
| Unique reflections | 17711 (1635) |
| Completeness (%) | 98.81 (93.25) |
| Wilson B-factor | 23.24 |
| Reflections used in refinement | 17699 (1630) |
| Reflections used for R-free | 888 (77) |
| R-work | 0.2370 (0.3426) |
| R-free | 0.2669 (0.3299) |
| Number of non-hydrogen atoms | 1375 |
| macromolecules | 1259 |
| ligands | 55 |
| solvent | 61 |
| Protein residues | 159 |
| RMS(bonds) | 0.008 |
| RMS(angles) | 0.97 |
| Ramachandran favored (%) | 98.73 |
| Ramachandran allowed (%) | 1.27 |
| Ramachandran outliers (%) | 0.00 |
| Rotamer outliers (%) | 0.00 |
| Clashscore | 13.16 |
| Average B-factor | 29.06 |
| macromolecules | 28.95 |
| ligands | 30.00 |
| solvent | 30.62 |

<sup>a</sup>Statistics for the highest-resolution shell are shown in parentheses.

**Figure 2.**
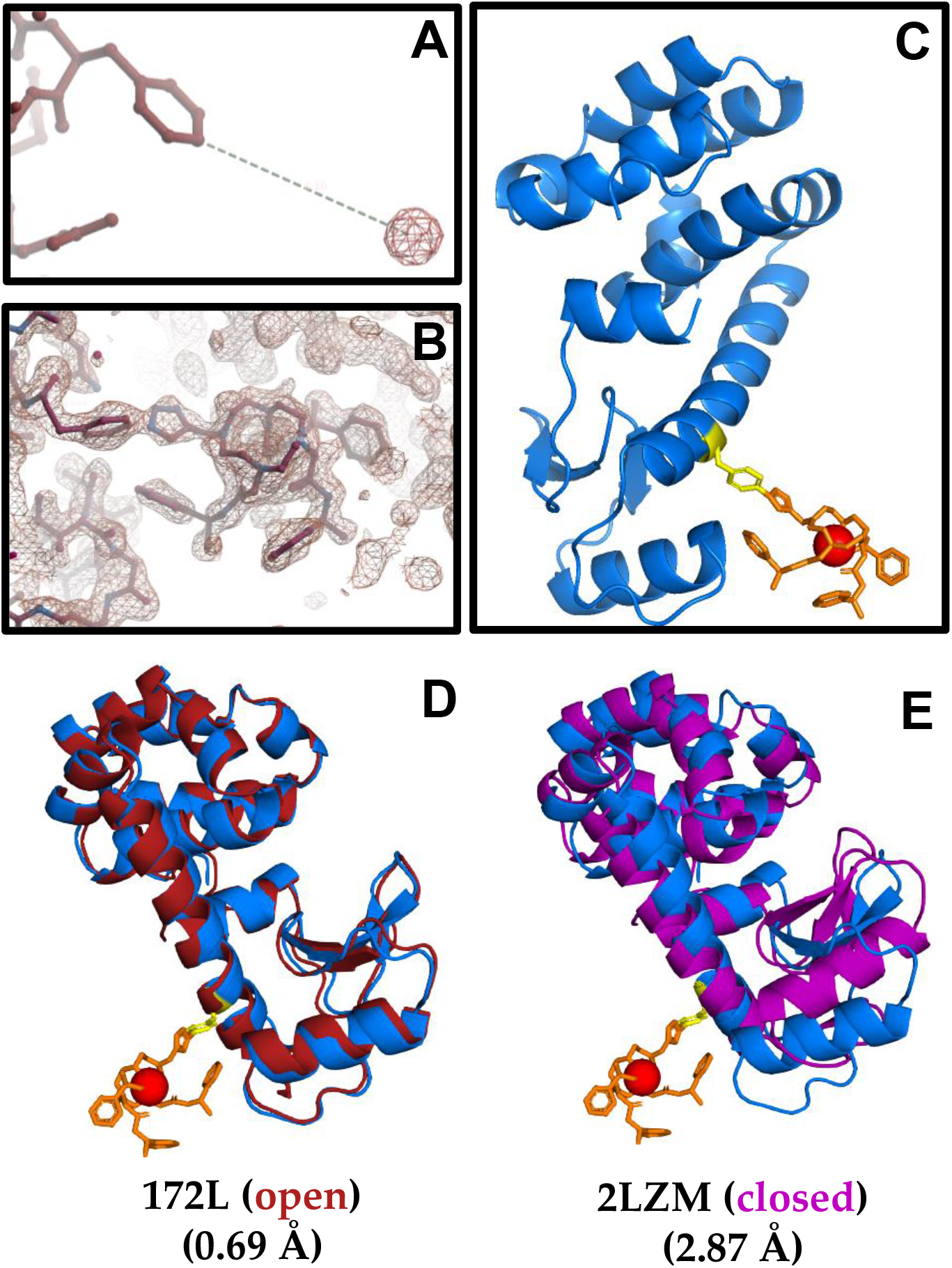
Crystal structure determination of lanthanide-labeled T4L. Structure determination and crystallographic analysis of C3-Tb^3+^-labeled protein shows (A) the Tb^3+^ ion (electron density shown in red) located 8.16 Å away from the CZ of the phenyl group at pAzF65 in T4L. (B) C3-Tb^3+^ was modeled into the electron density whose position could be easily interpreted. (C) The final model (blue) shows pAzF in yellow sticks, with the triazole ring formed from the azide-alkyne cycloaddition reaction. The three chiral pendants of the C3 tag (orange) participate in chelating the Tb^3+^ ion (shown as red sphere). Crystallographic evidence of the label conjugated to the protein confirms labeling with pAzf-C3-Tb^3+^ does not perturb native fold of the protein. T4L-pAzF-65-C3-Tb^3+^ crystallized in the open form with 0.690 Å RMSD compared to one of the most open forms observed in the PDB (172L), shown in red (D), and 2.867 Å RMSD relative to the wild type structure, which represents a closed form (2LZM) shown in purple (E). Detailed crystal parameters are given in Table 1.

### Crystallographic analysis of lanthanide-labeled protein shows T4L in an open conformation, and confirms that the tag does not perturb the structure of T4L

T4L-pAzF-65-C3-Tb^3+^ crystallized in the open configuration. This has been previously observed for only 10% of T4L mutants deposited in the PDB. When compared to available T4L structures in the PDB, our structure agrees best with one of the most open forms observed (PDB ID: 172L) at 0.69 Å RMSD (Figure 2D), and with the closed form observed for wild type (PDB ID:2LZM) at 2.87 Å RMSD (Figure 2E).

It was considered whether the open conformation could have been induced and/or perturbed by the presence of the C3 tag; however, this is unlikely as a similar open conformation has been observed under different crystallization conditions and in the absence of a tag (PDB ID: 172L). Hinge-bending in T4L, which governs its open vs. closed conformation transition, is suggested to be an intrinsic property of the molecule and is not an artifact due to mutation (Zhang et al., 1995). Furthermore, an EPR study by Mchaourab *et al*. demonstrated that an I3P mutation, which produced a large hinge-bending angle in the crystal, had no effect on the solution conformation (Mchaourab et al., 1997). This led to their conclusion that the hinge-bending motion in T4L is not the result of a mutation but is an integral part of T4L dynamics in solution.

### Comparison of crystallized pAzF-C3 conformation with theoretical dihedral angle preferences

Energy calculations on the M06/Lan12dz level of theory were run by systematically varying each of four side chain dihedral angles of the pAzF-C3 label to predict low-energy conformations for comparison to the observed conformation in the crystal structure. The calculated energies from these dihedral scans are shown in Figure 3. Similar results were obtained when using the BCL conformer generator method (Kothiwale et al., 2015; Mendenhall et al., 2021) to create low-energy conformations of the pAzF-C3 label by sampling rotamers from the crystallographic open database (COD) [see Supplementary Figure 1]. The following trends are observed: For χ_1_, the favorable angles are around 60°, 180°, and 300°. For χ2, the 90° and 270° are energetically most favorable. For χ_3_, the energy barriers are much lower (only ∼3 kcal/mol compared to ∼7 kcal/mol for χ_1_). Consequently, there are no distinct peaks in the distribution, but the rotamers are sampled more broadly. For χ_4_, the favorable regions are around 120° and 300°. Some “forbidden” regions (e.g. between 0 - 60°) are observed for χ_4_, where the energy is very high because of intra-molecular steric clashes.

**Figure 3.**
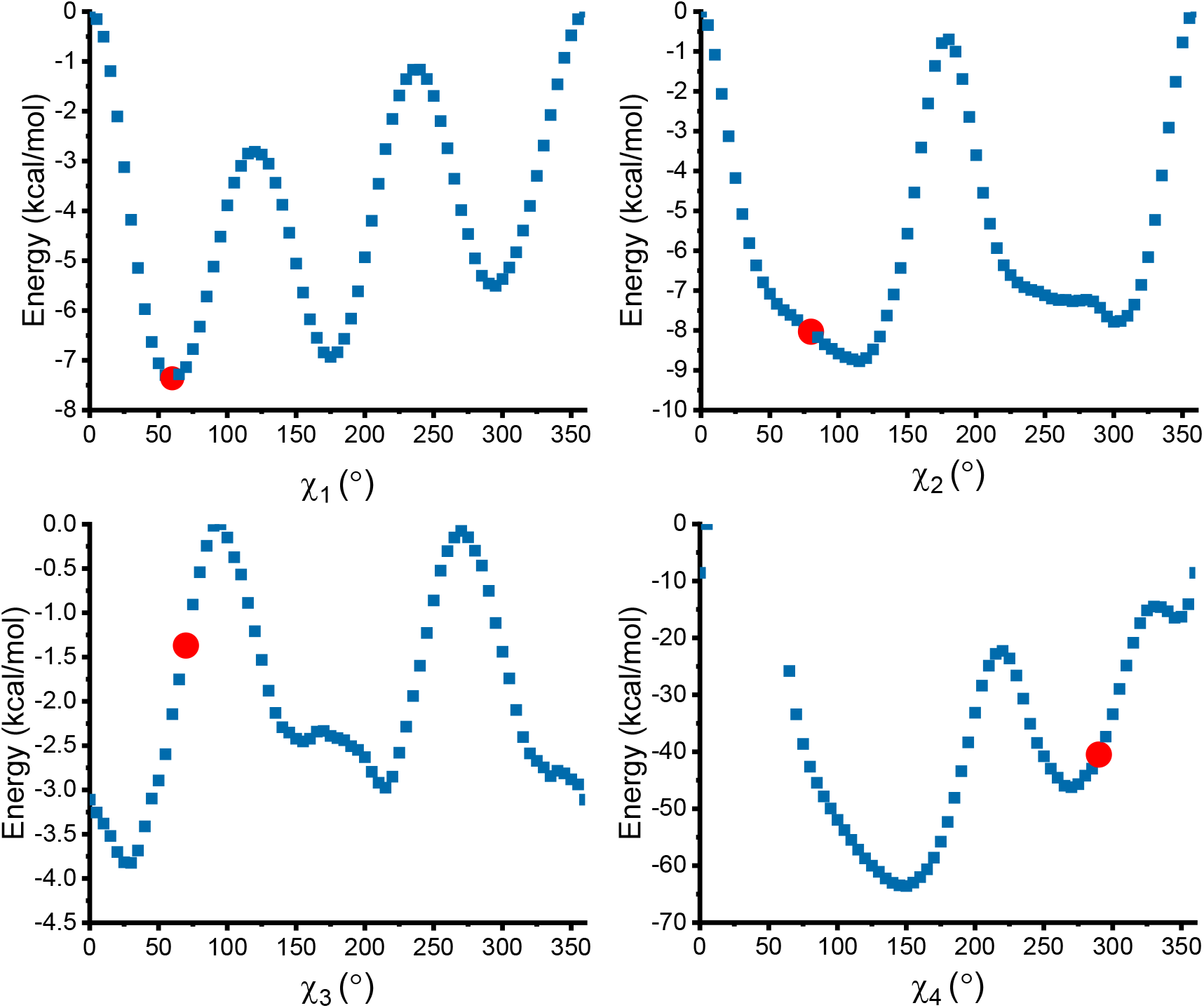
Theoretical predictions of pAzF-C3-lanthanide label conformations. The dihedral angle energy profiles for χ_1_ to χ_4_ determined by DFT. The chi angles observed in the crystal structure are indicated on each plot with a red circle.

The χ angles observed in the crystal structure are indicated on each plot in Figure 3: χ_1_: 59.5°; χ_2_: 78.3° χ_3_: 70.5° χ_4_: -67.4° (marked at 292.6°). χ_1_ and χ_2_ fall in the minimum energy well observed in the calculations; this shows the crystal structure adopted one of the lowest-energy conformations for χ_1_ and χ_2._ A low-energy conformation is also observed for χ_4_ in the energy well. χ_3_ is not observed at a minimum; however, as mentioned previously, the energy barriers are much lower for χ_3_. Thus, there appears to be less of a preference for χ_3_ which explains why the crystal structure does not adopt the lowest energy state.

### PCS measurements for use as paramagnetic structural restraints

2D [^1^H, ^15^N]-IPAP (Ding and Gronenborn, 2003; Ottiger et al., 1998) or 2D [^1^H, ^15^N]-HSQC NMR spectra of the four pAzF mutants labeled with paramagnetic lanthanides Yb^3+^, Tm^3+^, or Tb^3+^ were measured, and the C3-lanthanide label induced sizeable PCSs (Figure 4). PCSs are induced by each paramagnetic lanthanide ion (Yb^3+^, Tm^3+^, or Tb^3+^) as demonstrated for the selected resonances. Tm^3+^ resulted in PCSs with opposite sign to Tb^3+^, and different lanthanides yielded PCSs with different magnitudes, such that Tb^3+^ resulted in the largest PCSs. The observed ^1^H PCS values had magnitudes in the range for the following: T4L-pAzF65-C3-Yb^3+^ (−0.18 ppm to 0.61 ppm) (Supplementary Table 1), T4L-pAzF65-C3-Tm^3+^ (−0.49 ppm to 1.09 ppm) (Supplementary Table 2), and T4L-pAzF65-C3-Tb^3+^ (−1.54 ppm to 1.54 ppm) (Supplementary Table 3).

**Figure 4.**
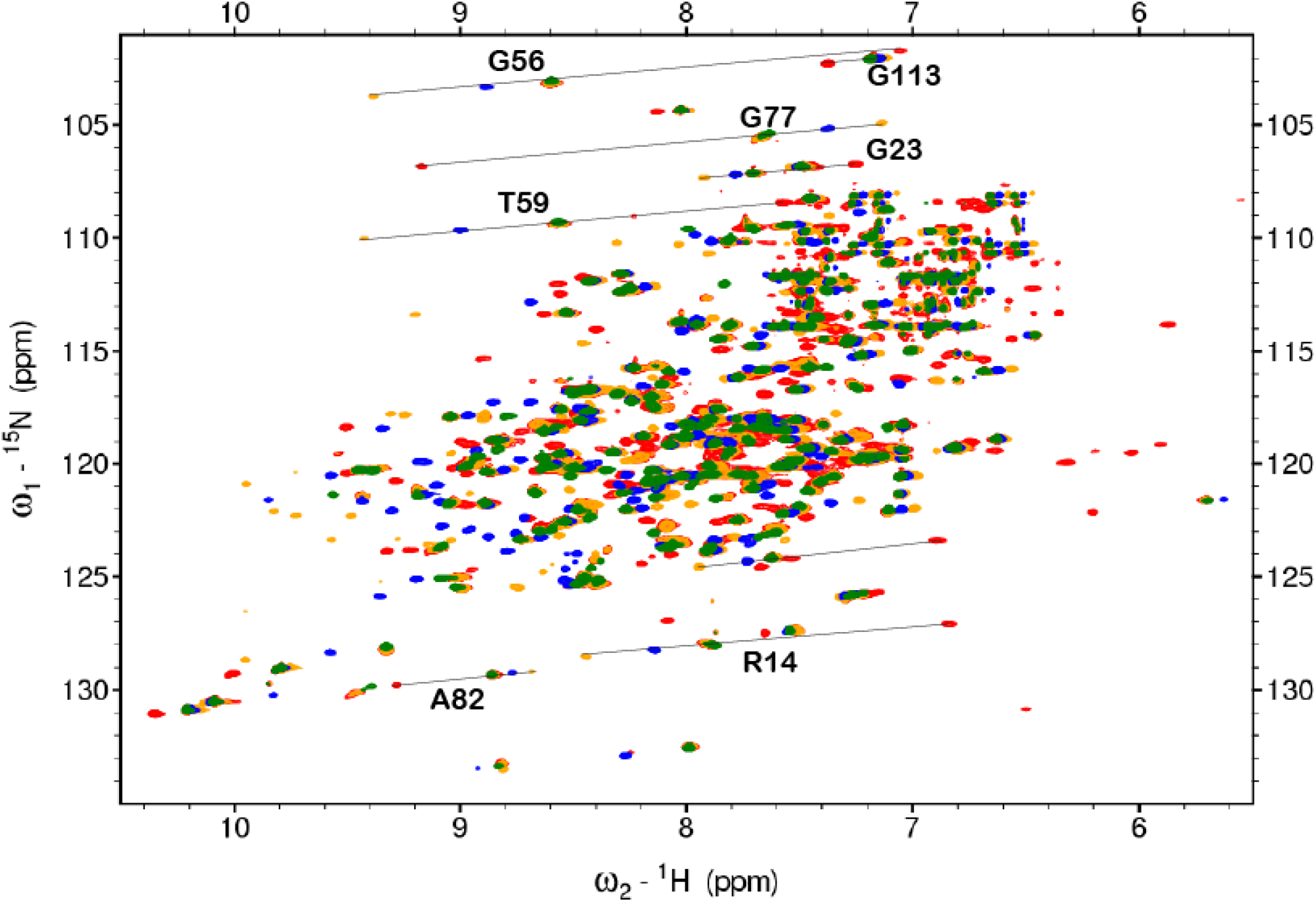
PCS dataset at position 65. Superimposition of 2D [^1^H-^15^N]-IPAP spectra of T4L-pAzF65 in complex with one diamagnetic C3-Y^3+^ (green), or paramagnetic C3-Yb^3+^ (blue), C3-Tm^3+^ (yellow), and C3-Tb^3+^ (red). Lines indicate a selection of observed PCSs. As seen for selected resonances, Yb^3+^ induces smaller PCSs, and Tb^3+^ induces the largest. Generally, Tm^3+^ induces PCSs with opposite sign to Tb^3+^. Spectra were recorded at 800 MHz ^1^H frequency and 298.0 K.

Overall, from the four mutants using the three paramagnetic lanthanide ions, PCS datasets for all three metals at position 65, two metals at position 119 and 131, and one metal at position 44 were obtained. PCSs were unambiguously assigned for backbone amides and backbone protons, and the tryptophan side chain indoles. A total of 345 PCSs was measured. For positions 44, 65, 119, and 131 18, 37, 25, and 18 PCSs were determined, respectively. Significant PCSs were generated for the sample labeled at position 65 in the presence of all three paramagnetic lanthanides. Fewer PCSs could be derived for Tm^3+^ or Tb^3+^ because signals were broadened beyond detection. PCSs generated at positions 119 (Supplementary Figure 2) and 131 (Supplementary Figure 3) were measured in the presence of only Tm^3+^ or Tb^3+^; PCSs in the presence of Yb^3+^ were too small to be observed. PCSs at position 44 (Supplementary Figure 4) were only measured for Tm^3+^; PCSs in the presence of Yb^3+^ were too small to be observed, while in the presence of Tb^3+^, PCSs were broadened beyond detection. The assigned PCSs for these mutants are reported in Supplementary Tables 4 – 8.

Along with the analysis of the 2D NMR spectra, crystallographic evidence and circular dichroism spectra showed that the predominantly α-helical structure typical for T4L remained unchanged (Supplementary Figure 6), and confirmed the structural integrity of the C3-pAzF-labeled mutants.

### PCS data fit better to a more open T4L conformation in solution

For a protein with known structure, the anisotropic part of the magnetic susceptibility tensor (Δχ) can be determined by simultaneous fitting of PCS values. The experimental PCS values measured for the T4L mutants pAzF65, pAzF119, pAzF131, and pAzF44 were used to fit Δχ tensors to the closed conformation of T4L, PDB ID 2LZM, or the open conformation (i.e. the structure determined in this work), using the program Paramagpy (Orton, 2020) (Figure 5 – top panel; Supplementary Figure 2 - 4). Δχ tensors can be visualized as isosurfaces. Example isosurfaces are shown for position 65 with Yb^3+^ and Tb^3+^ (Figure 5 – bottom panel).

**Figure 5.**
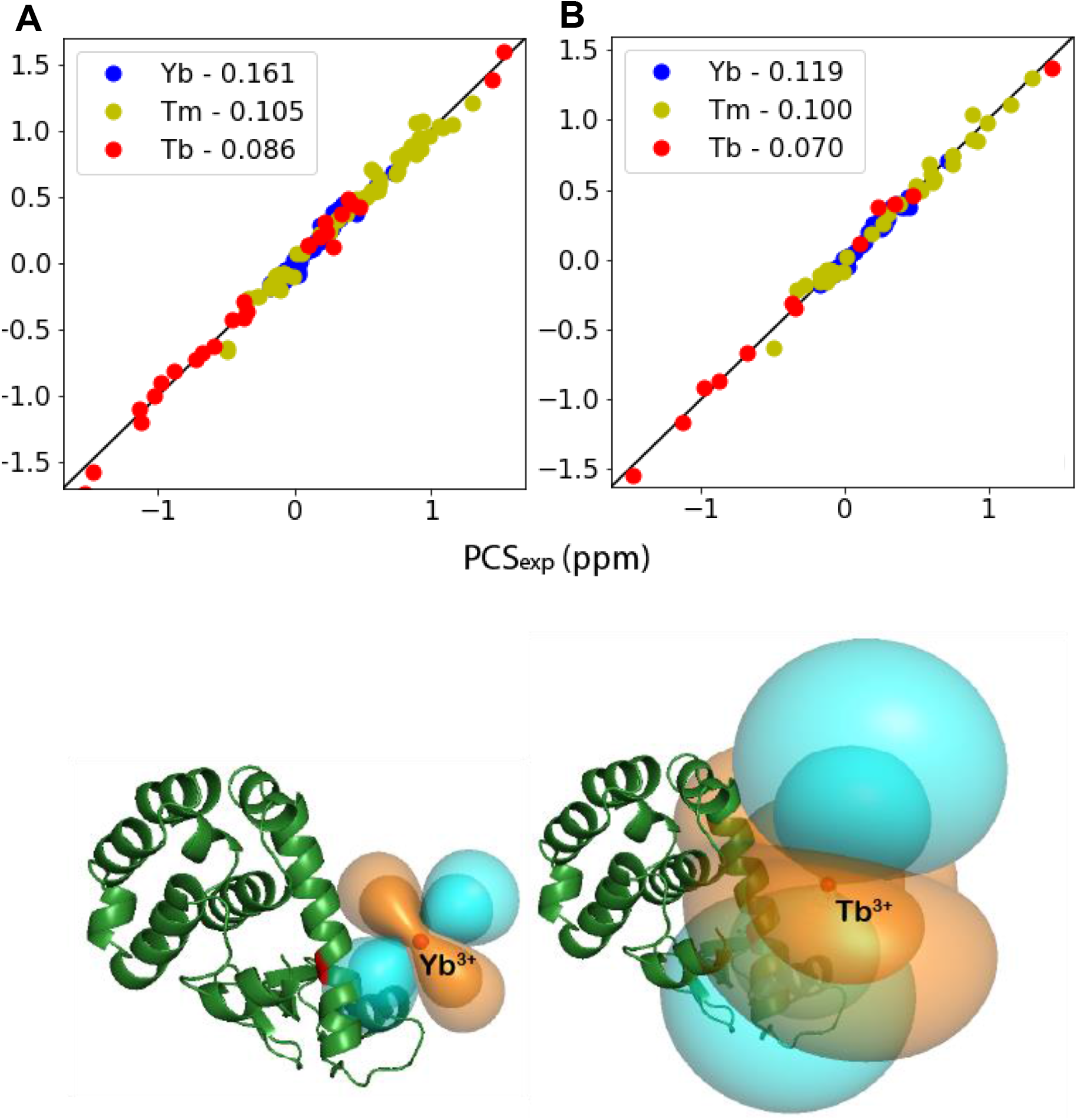
Correlation between observed and calculated PCS values for T4L-pAzF65. Top panel: The PCS data for both backbone amide protons and nitrogens obtained with Yb^3+^ (blue), Tm^3+^ (yellow), and Tb^3+^ (red) coordinated to the C3 label were fitted globally to (A) the closed structure of T4L (PDB ID: 2LZM) and (B) the open structure of T4L (determined in this work). The PCS Q-factors between the observed and calculated values are given. Bottom panel: The positions of Yb^3+^ and Tb^3+^ are indicated with spheres, and the associated PCS isosurfaces are shown - cyan (+0.5 ppm and +2.5 ppm) and orange (−0.5 ppm and -2.5 ppm).

For each mutant, PCSs were fit simultaneously for all lanthanides at a common metal ion position, except for S44pAzF, which was fit using only one lanthanide ion. The metal position, considering the rigidity of the tag, should change very little between the different lanthanides because of their similar chemical properties and because they are shielded by the chelator. Even though the metal coordinates (Mx, My, Mz) remain the same, alpha, beta, gamma, χ-axial, and χ-rhombic, are still metal-specific as seen in the Δχ tensor parameters reported for each site (Table 2). Tensor fit quality was assessed by PCS Q-factor (Chen et al., 2016). The Δχ tensors determined with the C3 label at each position are reported. The back-calculated PCS values are in good agreement with the observed PCSs, with PCS Q-factors mostly below 0.2. Specifically, at position 65, Q-factors ranged from 0.086 (Tb^3+^) to 0.119 (Yb^3+^); position 119: 0.048 (Tb^3+^) to 0.065 (Tm^3+^); position: 131: 0.054 (Tb^3+^) to 0.098 (Tm^3+^); position 44: 0.140 (Tm^3+^). The low Q-factors indicate rigidity of the C3 tag, and reliability of the PCS measurements in agreement with the structural models. This is important because high flexibility of a paramagnetic label can convolute the measurement or prediction of PCSs close to the paramagnetic center.

**Table 2.** The Δχ-tensor parameters determined for the C3-lanthanide label.

| **Ln<sup>3+</sup>** | **$\Delta\chi_{ax}$**<br>**(10<sup>-32</sup> m<sup>3</sup>)** | **$\Delta\chi_{rh}$**<br>**(10<sup>-32</sup> m<sup>3</sup>)** | **x (Å)** | **y (Å)** | **z (Å)** | **$\alpha$ (°)<sup>1</sup>** | **$\beta$ (°)** | **$\gamma$ (°)** |
| --- | --- | --- | --- | --- | --- | --- | --- | --- |
| **65-Yb<sup>3+ 2</sup>** | -6.05 | -1.95 | 34.0 | 55.3 | 12.3 | 37.4 | 97.1 | 30.2 |
| **65-Tm<sup>3+</sup>** | -14.35 | -5.45 | 34.0 | 55.3 | 12.3 | 30.7 | 96.1 | 9.2 |
| **65-Tb<sup>3+</sup>** | 30.87 | 9.89 | 34.0 | 55.3 | 12.3 | 31.0 | 89.8 | 10.6 |
| **119-Tm<sup>3+</sup>** | -27.54 | -4.61 | 49.74 | 60.81 | 49.97 | 109.8 | 127.4 | 117.1 |
| **119-Tb<sup>3+</sup>** | 20.1 | 6.25 | 49.74 | 60.81 | 49.97 | 98.5 | 134.3 | 94.6 |
| **131-Tm<sup>3+</sup>** | -20.95 | -6.68 | 43.8 | 37.7 | 54.9 | 64.0 | 60.0 | 59.0 |
| **131-Tb<sup>3+</sup>** | 23.96 | 7.30 | 43.8 | 37.7 | 54.9 | 89.5 | 54.4 | 76.9 |
| **44-Tm<sup>3+</sup>** | -16.13 | -3.99 | 16.0 | 52.1 | 11.9 | 58.0 | 111.0 | 112.0 |

<sup>1</sup> The Euler angles ( $\alpha$ , $\beta$ , $\gamma$ ) for the PCS tensors are in ZYZ convention with units of degrees.
<sup>2</sup> Tag coordinates for position 65 were determined from the crystal structure of T4L-65-pAzF-C3-Tb<sup>3+</sup> (described in this paper).

Although PCS Q-factors were low for both the open and closed structures of T4L (Figure 5B, and Supplementary Figure 2 - 4) better agreement was observed when using the more open conformation of T4L and the measured PCSs, suggesting the T4L conformation observed is more open in solution. This is supported by previous studies of T4L conformation in solution which demonstrated T4L conformation in solution is likely more open than suggested by the observed crystal forms of T4L, including the wild type. It is not surprising that we observed low PCS Q-factors for both closed and open models because T4L displays a wide range of conformations in the N-terminal region, owing to the hinge-bending motion. To confirm this, PCS measurements for residues in the flexible N-terminal were fit to T4L structure (open or closed). The following are the average Q-factors calculated using PCSs measured for open or closed conformation: pAzF65 (0.05 - open; 0.06 – closed), pAzF119 (0.09 – open; 0.12 - closed), pAzF131 (0.12 – open; 0.17 – closed) [Supplementary Table 9]. The PCSs from the N-terminal region are in better agreement with the open conformation. Thus, the measured PCSs captured the more structured α-helical region in both open and closed forms, yielding fits with similar good PCS Q-factors. This suggests that the slight improvement in Q-factors calculated for the open structure takes into account the flexible N-terminal domain, and in this case, the better fit can be attributed to the measured PCSs capturing T4L in a more open state.

### De novo folding of T4L with Rosetta using PCSs measured at multiple sites

To assess the performance of computational structure prediction using the sparse PCS datasets, the structure of T4L was re-determined *de novo* (from sequence only without any homology modeling) in the computational modeling package, Rosetta, using only PCS information. The goal was to analyze 1) the improvement of structure prediction (determined by the number of models generated that were closer to the native structure), and 2) the ability of PCSs to capture the structure of the more flexible domain, thereby allowing the observation of alternative conformations. More importantly, we were interested in the level of improvement that would be observed when using PCSs measured at multiple sites as compared to using PCSs from a single site.

10,000 models each were generated with RosettaNMR in the presence or absence of PCS data. Rosetta performance with PCSs and other paramagnetic data have been discussed and benchmarked previously (Kuenze, 2019; Schmitz et al., 2012) and thus is not discussed in detail in this paper. The focus here is on modeling of T4L using sparse PCSs solely obtained from the experimental protocol employed in this work. To assess the contribution of PCSs obtained from multiple sites, calculations were run using (1) no PCS data, (2) PCSs measured at one site (position 65), (3) PCSs measured at two sites (positions 65 and 131), (4) PCSs measured at three sites (positions 65, 131, and 119), and finally (4) PCSs measured at four sites (positions 65, 131, 119, and 44). All computations used the same fragment set (which excluded all fragments from structures of T4L in the PDB). For running *de novo* structure prediction, the AbRelax protocol in RosettaNMR was used. For the folding stage, a weight of the PCS score that was optimized against the score3 energy was employed. For final model scoring and selection the PCS weight was adjusted relative to the total ref2015 energy.

### Improved conformational sampling by RosettaNMR with measured PCSs

PCSs bias sampling toward the native structure (Figure 6). The improvement in model quality was pronounced when PCSs obtained from different sites were included in the calculation. Specifically, the number of models (out of 10,000) generated that were less than 7 Å from the reference structure was: 2 (in the absence of PCSs, with lowest RMSD of 6.6 Å), 138 (PCSs from one site (pAzF65), with lowest RMSD of 3.5 Å), 157 (two sites (pAzF65 + pAzF131), lowest RMSD of 2.8 Å), 268 (three sites (pAzF65 + pAzF131 + pAzF119), lowest RMSD of 2.9 Å), and 267 (four sites (pAzF65 + pAzF131 + pAzF119 + pAzF44); lowest RMSD of 2.9 Å). Final models selected for each run are shown in Figure 7. For the trial with no PCS restraints, the top scoring model (Rosetta score) had an RMSD of 11.46 Å to native T4L. When PCSs from one site were used, the top-scoring model (Rosetta + PCS score) had an RMSD of 7.60 Å to native T4L. For PCSs from two, three, and four sites, the top-scoring (Rosetta + PCS) models had RMSDs of 8.52 Å, 7.58 Å, and 7.64 Å, respectively.

**Figure 6.**
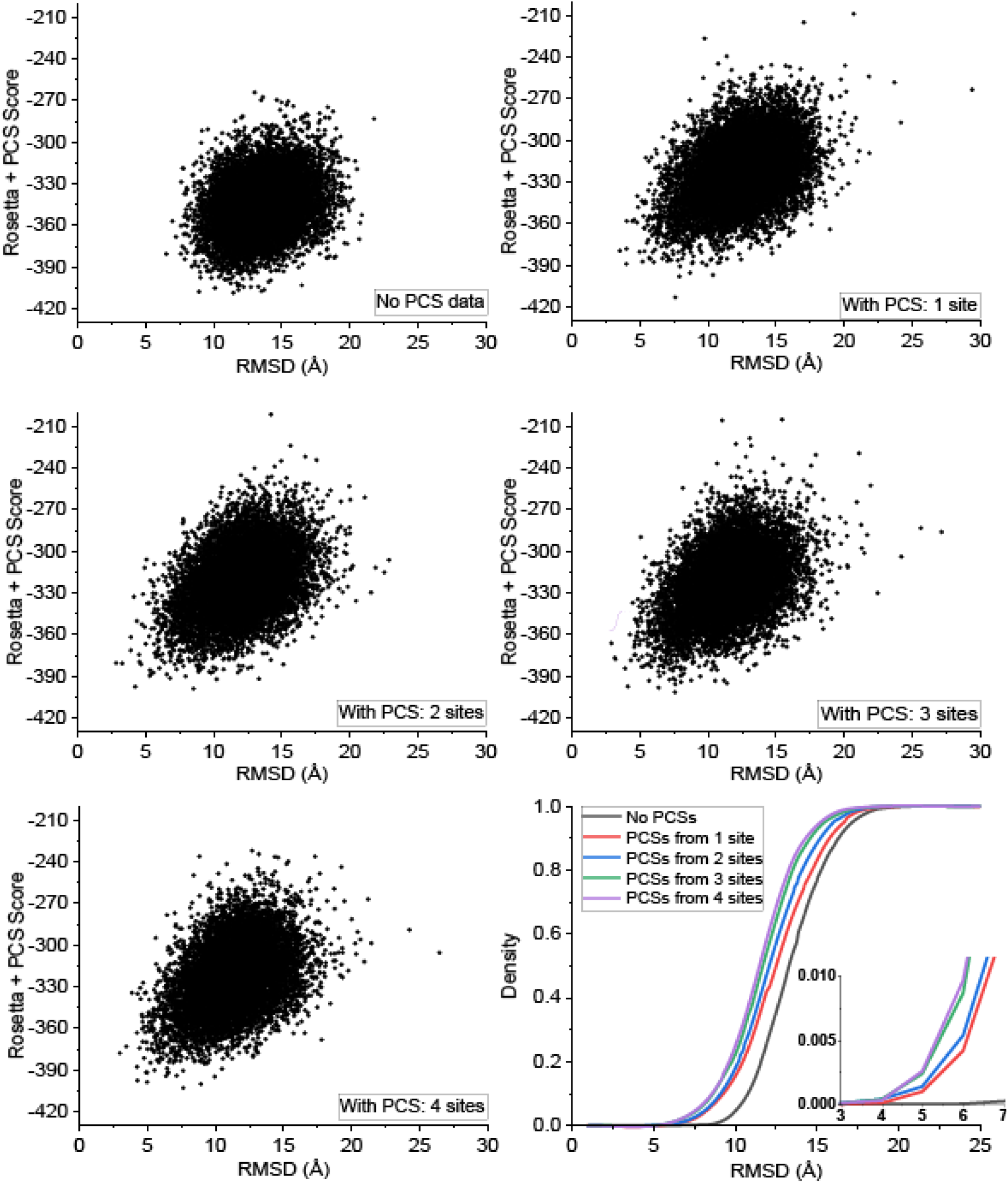
Improved conformational sampling by Rosetta with PCSs. The score-vs-RMSD plots show the energy landscape generated by Rosetta using no PCS data or with PCS data measured from up to four sites. The density plot at the lower right corner shows marked improvement in sampling of low-RMSD models when PCSs generated at multiple sites were used. The results suggest that ROSETTA with PCSs can generate more accurate models and identify the correct protein fold.

**Figure 7.**
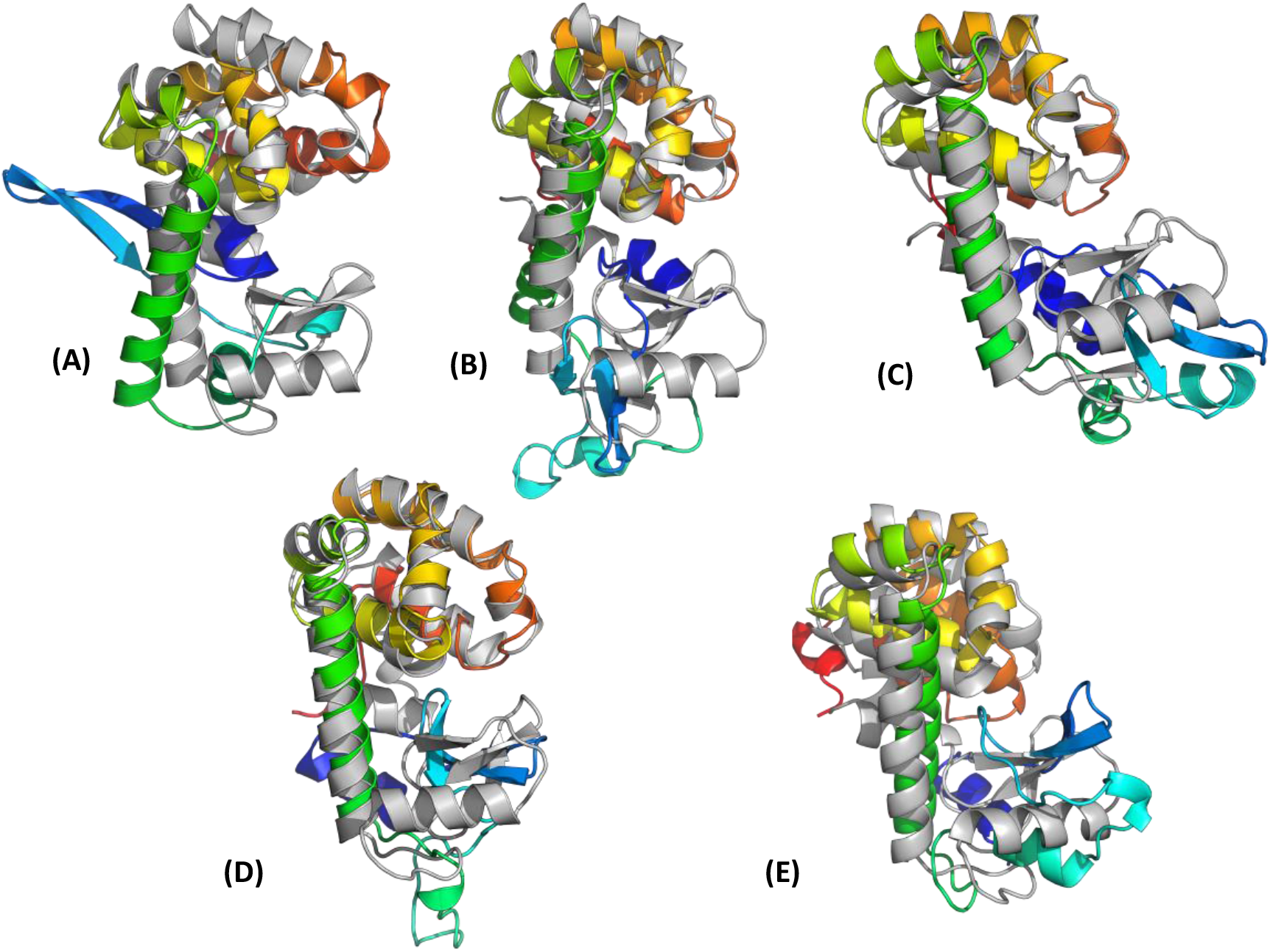
T4L best-scoring models obtained by Rosetta structure calculations using sparse PCS restraints from multiple labeling sites. Displayed are the final models selected based on lowest score from each run (rainbow colored) superimposed to T4L native structure (gray). Superimposition of the lowest scoring (ROSETTA + PCS score) model in runs with (A) no PCS restraints - RMSD to native: 11.46 Å; (B) PCS restraints from one site – RMSD: 7.60 Å; (C) PCS restraints from two sites – RMSD: 8.52 Å; (D) PCS restraints from three sites – RMSD: 7.58 Å ; (E) PCS restraints from four sites – RMSD: 7.64 Å.

The lowest RMSD model obtained from each run is displayed in Figure 8. For structure calculations with no PCSs, lowest RMSD to native T4L is 6.51 Å, and for calculations with PCSs from one, two, three, or four sites, lowest RMSDs to native T4L are 3.53 Å, 2.83 Å, 2.91 Å, and 2.97 Å, respectively. Significant improvement is observed in modeling the full-length protein when PCSs from at least one site are used. Using PCSs from multiple sites resulted in an improved structural model.

**Figure 8.**
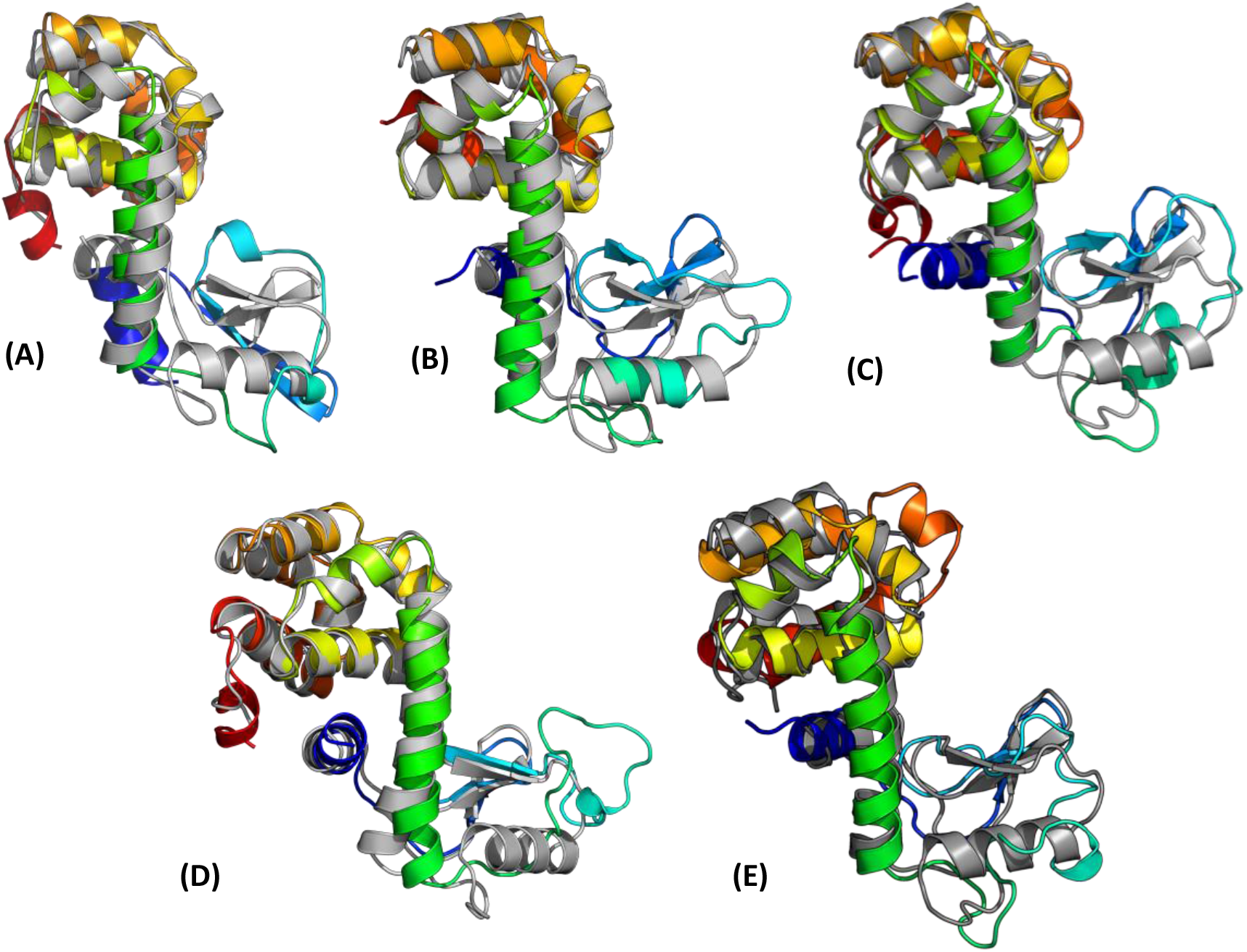
T4L lowest-RMSD models obtained by ROSETTA structure calculations using sparse PCS restraints from multiple labeling sites. Displayed are the lowest-RMSD models selected from each run (rainbow colored) superimposed to T4L native structure (gray). Superimposition of the lowest RMSD model in runs with (A) no PCS restraints - RMSD to native: 6.51 Å; (B) PCS restraints from one site – RMSD: 3.53 Å; (C) PCS restraints from two sites – RMSD: 2.83 Å; (D) PCS restraints from three sites – RMSD: 2.91 Å ; (E) PCS restraints from four sites – RMSD: 2.97 Å. ROSETTA generates models closer to the native when PCSs are included in modeling. As seen for these models, there is marked improvement in RMSD to the native not only in the structured α-helical regions, but also in the more flexible regions.

The results here show that sparse datasets from a single site (with experimentally-determined tag position and orientation) were powerful enough to significantly increase the number of low-energy models close to the native structure sampled by Rosetta (from two to 138 models below 7 Å). This means the measured PCSs provide reliable restraints that correctly guide Rosetta in *de novo* structure determination. Furthermore, datasets from PCSs measured at multiple positions improve not only sampling of models close to the native structure, but there is a marked improvement for the modeling in the structured α-helical and flexible regions. Without PCSs, no convergence is observed in the ensemble of the top 10 models (lowest-scoring) generated of full-length T4L in either the structured or flexible domains (Supplementary Figure 7). There is improved convergence to the native structure with restraints from only one site. Nonetheless, even though the structured region is sampled with low RMSD more frequently, the more flexible domain still has increased RMSD values relative to the native structure (Figure 7B and Figure 8B). Calculations including PCSs at three or four sites sampled the structured regions with low RMSD more frequently and resulted in models that superimposed very well to the flexible domain on the native structure. Combining PCS score with Rosetta score (or using PCS score only) led to generating and identifying models with RMSD closest to the native (Figure 9). Furthermore, modeling with PCSs from multiple sites showed a marked improvement. These results confirm that attaching paramagnetic tags at multiple locations to increase coverage of the protein results in high quality distance and orientational PCS restraints that better guide computational structure modeling protocols.

**Figure 9.**
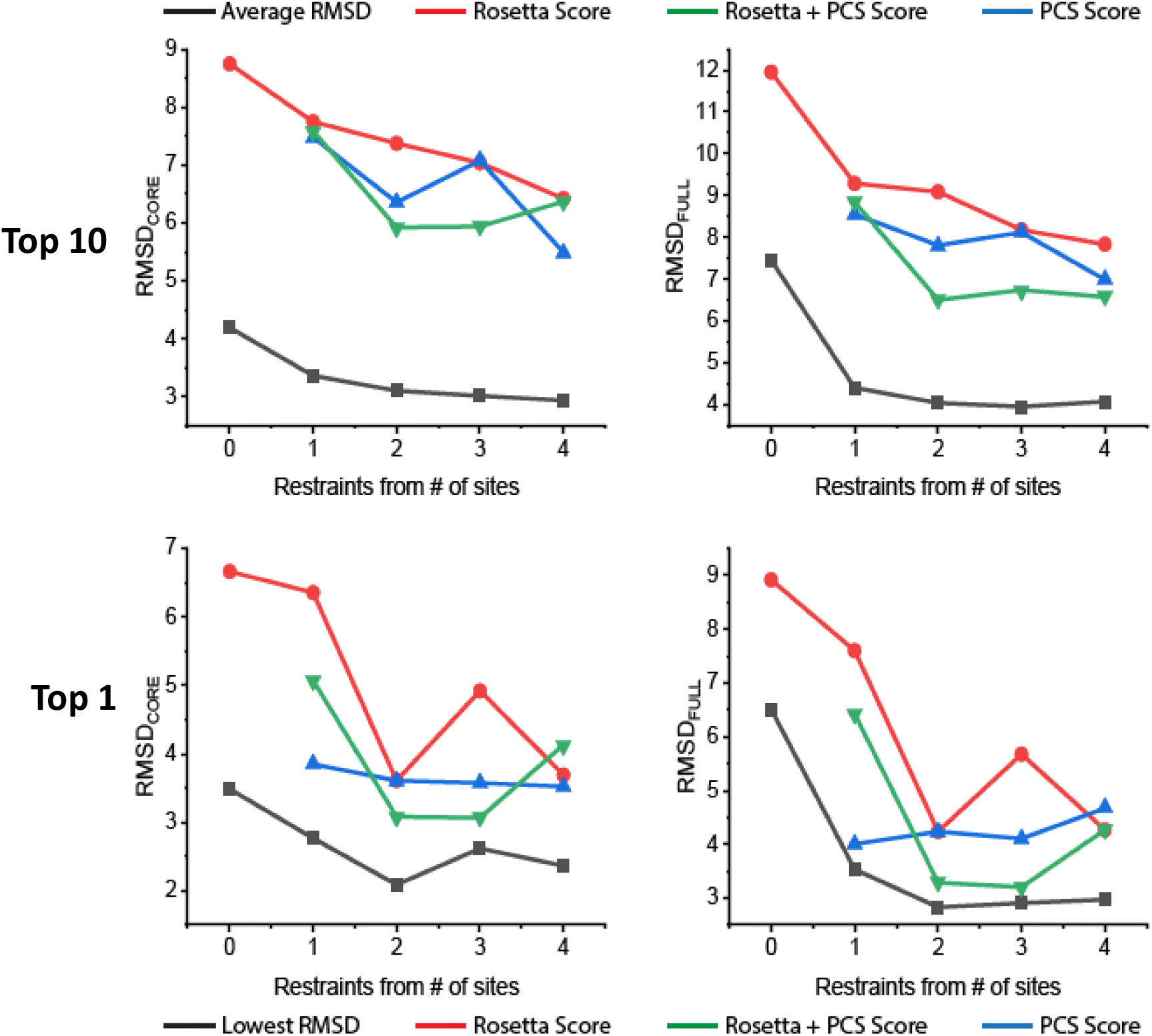
Analysis of ensembles of final models selected in Rosetta. Top panel: Analysis of top 10 best-scoring models generated in Rosetta. Shown in black is the average of all pairwise RMSDs in the ensemble for each run using PCSs from a different number of sites. Average RMSD of ensemble selected by ROSETTA score is in red, by ROSETTA + PCS score in green, by PCS score only, in blue. The left panel shows measurements for the core α-helical region and in the right panel, the full-length protein. In both cases, combining PCS score with ROSETTA score improved model selection significantly, while PCS scores identified models closest to the native. Bottom panel: The analysis was repeated for the lowest RMSD models generated in each run. Black line shows lowest RMSD model from each run. Red line shows lowest RMSD model when selected by ROSETTA score, green line for selection by ROSETTA + PCS score, and blue line for selection by PCS score only. In both cases, the combined ROSETTA+PCS score or PCS score was better at identifying models with low RMSD to the native structure.

## Conclusions

PCSs are a rich source of structural information through the interrogation of individual atoms of a biomolecule. In recent years, there has been increased interest in lanthanide labels for PCS measurement. In this work, limited PCS datasets were obtained from multiple sites in the bacteriophage model protein T4L and used to demonstrate the reliability of the lanthanide-induced PCSs for computational modeling techniques. The datasets led to successful re-determination of the three-dimensional structure of T4L and assessed conformations of T4L in solution using PCSs only. These results demonstrate the power of lanthanide-induced PCSs from multiple sites for structure determination and conformational analysis of flexible systems. The PCSs measured successfully guide the folding of T4L and capture the conformational dynamics of T4L in solution.

We present a complete experimental and computational pipeline for protein structure modeling leveraging the attachment of a paramagnetic tag via the ncAA, pAzF. In addition to reporting the first experimental structure that showcases both the tag location and orientation, we show that PCSs generated in this manner at multiple labeling sites significantly improve *de novo* protein structure prediction. This may prove powerful when coupled with computational modeling techniques to study protein structure, conformational dynamics, and interactions between biomolecular complexes. Advances in computational structure prediction (e.g. RoseTTAfold and Alphafold2), presents an exciting approach for experimentalists to provide validation and information to help interpret the biological relevance and biophysical significance of computational models. The ease of measuring PCSs and the versatility of lanthanides for this purpose could render PCSs one of the most useful types of structural restraints for studying the conformation and dynamics of flexible systems by solution NMR. The pipeline described here is equally applicable to the study of larger protein complexes in solution, including integral membrane proteins.

## STAR Methods

### Protein Expression and Purification

Site-specific incorporation of para-Azido-phenylalanine (pAzF) was done using the published pEVOL vector with the aminoacyl-tRNA synthetase for pAzF (pAzF-RS)(Young et al., 2010). Amber stop codons (UAG) were introduced in the T4-lysozyme (T4L) sequence, complemented by orthogonal translation machinery provided by the pEVOL plasmid. Plasmids coding for the T4L mutants and pEVOL-pAzF plasmid encoding pAzF-tRNA synthetase and tRNA were co-transformed into *E. coli* BL21(DE3) cells and grown at 37°C in the presence of 100 μg/mL ampicillin and 33 μg/mL chloramphenicol.

T4L mutants were expressed with an amber stop codon to replace residue Ser44, Lys56, Thr109, Arg1199 or Val131 with pAzF. 10mL of an overnight culture was spun down, resuspended in 10mL of ultrapure water and used to inoculate 500 mL minimal media for ^15^N-labeling. Protein was produced by growing cells in M9 medium containing 0.5 g/L ^15^NH_4_Cl and 0.02% arabinose. After growing to an OD_600_ of 0.7 – 1.0, protein expression was induced with the addition of 1 mM pAzF and 1 mM isopropyl-β-D-thiogalactopyranoside (IPTG), and incubated with shaking at 37°C for 1.5 - 2 hours. The cultures were harvested by centrifugation, and the pellets were resuspended in lysis buffer (20 mM MES, 20 mM TRIS, 0.01% sodium azide, at pH 6.5 containing protease tablet (Sigma Aldrich), upon which the cells were lysed by sonication. The cell lysates were centrifuged for 20 min at 20,000 rpm, and supernatant was loaded onto a Resource S ion exchange (IEX) column (GE Healthcare Life Sciences, USA) and the protein was eluted with IEX buffer (20 mM MES, 20 mM TRIS, 0.01% sodium azide, at pH 6.5, supplemented with 1M NaCl). The fractions were analyzed by 10% SDS-PAGE, and the purified samples were concentrated using an Amicon ultrafiltration centrifugal tube with a molecular weight cutoff (MWCO) of 10 kDa.

### Ligation Reactions

Solutions of T4L containing the pAzF residue in 50 mM HEPES buffer, pH 7.5, were added to solutions of C3 label (prepared at 10 mM in water, by heating to dissolve, and then filtering) followed by addition of CuSO_4_ and copper-binding ligand BTTAA (2-(4-((bis[(1-tert-butyl-1H-1,2,3-triazol-4-yl)methyl]-amino)methyl)-1H-1,2,3-triazol-1-yl]acetic acid) (purchased from the Albert Einstein College of Medicine of Yeshiva University), and sodium ascorbate to a total reaction volume of 5 mL. The final concentrations were 0.5 mM protein, 1 mM C3 tag, 0.2 mM CuSO4, 1 mM BTTAA, and 5 mM sodium ascorbate. 5 mM aminoguanidine was added to the reaction buffer to prevent protein aggregation caused by byproducts from ascorbate oxidation. All ligation reactions were performed in a glovebox under N_2_ atmosphere at room temperature with gentle rotating for 16 – 18 h. The ligation reaction was terminated by the addition of 5 mM EDTA and standing for 30 min in air (Loh et al., 2013). The reactions were desalted using a Zeba spin desalting column (Thermofisher Scientific), and concentrated by ultrafiltration using an Amicon centrifugal filter (10,000 MWCO).

### NMR Spectroscopy

All NMR experiments were performed at 25°C on an 800 MHz Bruker Avance NMR spectrometer equipped with a cryoprobe. 3D spectra (CBCACONH, HNCA, HNCACB, HNCACO, HNCO, AND HNCOCA) were recorded of T4L to confirm resonance assignments using standard triple resonance techniques. PCSs were measured in the ^1^H and ^15^N dimension of ^15^N-HSQC (or ^15^N-IPAP) spectra. The spectra were recorded of 0.2 mM solutions of T4L mutants in complex with one equivalent of diamagnetic Y^3+^ or paramagnetic Yb^3+^, Tb^3+^, or Tm^3+^.

### Determination of the Δχ tensors

The experimental PCS values measured for the T4L mutants were used to fit magnetic susceptibility anisotropy tensors (Δχ) to the T4L crystal structure (PDB ID 2LZM) or the crystal structure determined in this study, using the program Paramagpy (Orton, 2020). Δχ tensors were determined using the ^1^H and ^15^N PCSs of the backbone amides. Tensor fit quality was assessed by Q-factor calculated as (Chen et al., 2016):

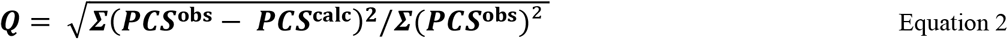

with PCS^obs^ being observed PCSs and PCS^calc^ being the back-calculated PCS values. The Δχ tensors determined with the C3 label at each position are reported in Table 2.

### Circular Dichroism (CD) experiments

CD was performed on a Jasco810 spectropolarimeter at room temperature. The sample concentrations were 12–16 μM in buffer consisting of 10 mM MES at pH 6.5. The path length of the optical cell was 0.5 mm, and the scan rate used was 50 nm/min with a response time of 1 s. Each spectrum shown is an average of nine scans.

### Crystallization and data collection

Initial crystallization conditions were identified using the vapor-diffusion technique in sitting drops in a high-throughput crystallization screen. Promising hits were optimized by mixing 1 μL or 2 μL of lanthanide-labeled T4L (20 mg/mL) with 2 μL reservoir solution and equilibrated against reservoir solution containing 0.2 M ammonium phosphate dibasic with 20% w/v PEG-3350. Crystals were grown at 21°C. Crystals appeared within two weeks. Crystals were mounted and flash-cooled in liquid nitrogen. Diffraction data were collected to a resolution limit of 0.9 Å. Full datasets with an interval of 0.5° were remotely collected at the LS-CAT beamline (Life Sciences Collaborative Access Team beamline 21-ID-G at the Advanced Photon Source, Argonne National Laboratory). All datasets were processed with HKL2000. A summary of the data statistics is given in Table 2.

### Structure determination, refinement, and analysis

The structure was resolved by molecular replacement in PHENIX using PDB 2LZM as a search model. Subsequent rounds of manual building using Coot and refinement using phenix.refine allowed complete model building and revealed clear density for the incorporated pAzF-C3-Tb^3+^ at position 65. The final model yielded crystallographic R-factors of 0.237/0.267 (R_work_/R_free_). The models were validated using MolProbity. Evaluation of the Ramachandran plot showed all residues in allowed regions (98.73 % in favored regions). All figures were prepared using Coot, Raster3D, and PyMOL. The data are deposited in the Protein Data Bank with PDB code 7U4X.

### Theoretical pAzF-C3 tag conformation calculations

To calculate the distribution of rotameric states sampled for the individual chi angles, 2000 conformers were generated of the pAzF-C3 spin label using the BCL (the Biochemical Library) conformer generator developed in the Meiler group. In addition, quantum mechanical (QM) calculations of the dihedral energy of chi angles χ1 to χ4 for the pAzF-C3 spin label were run using DFT with the M06 functional and Lanl2dz basis set. This includes atoms up to lanthanum (La), which was used as the metal in the C3 tag for calculations. pAzF-C3-La was capped with ACE and NME capping groups. Each of the four chi angles was scanned in 5° steps and the single point energy for the given conformation was calculated with Gaussian 09. The energy unit in Atomic Units (A.U.) or Hartree was converted to kcal/mol.

### Rosetta calculations

Calculations in Rosetta were performed as previously described (Kuenze, 2019). *De novo* structure prediction was completed using Rosetta’s fragment assembly protocol followed by all-atom refinement via FastRelax. Secondary structure information was employed for fragment selection, and PCS data was used for low-resolution fragment assembly, high-resolution refinement, model scoring and final model selection.

The fragment search was conducted with the Rosetta3 fragment picker application. A fragment library was generated for full length T4L using only sequence-based secondary structure, which was the consensus of two prediction methods, PSIPRED and Jufo9D. Homologous proteins present in the database were excluded according to a sequence similarity criterion (PSI-BLAST E-value < 0.5). To guide fragment assembly with PCSs, the score function at each of the four assembly stages was supplemented with the score term for the PCS restraint, using a weighting factor W_PCS_ optimized against Rosetta’s score3 energy function. After high-resolution refinement, models were rescored with the Rosetta all-atom energy function, combined with the PCS score term.

For each combination of PCSs 10,000 all-atom models were generated, and computations were run on the Vanderbilt University local ACCRE cluster. Rosetta models were assessed by calculating model RMSD to the native structure. Final models were selected based on RMSD to reference, combined Rosetta and PCS scores, and convergence.

## Acknowledgements

This work was supported by an NIH RO1 (Grant #: RO1 GM 0804-03). E.O. was supported by fellowships from the National Institutes of Health (Grant number: 5 F31 GM133134-02) and an institutional training grant (Molecular Biophysics Training Program) (Grant #: 5 T32 GM 8320-09). The work was conducted using the resources of the biomolecular crystallography facility, the biomolecular NMR facility, the synthesis core, and the resources of the Advanced Computing Center for Research and Education (ACCRE) at Vanderbilt University (VU). We would like to thank Dr. Markus Voehler of the NMR facility at VU, Dr. Kwangho Kim of the synthesis core at VU, and members of the RosettaCommons for discussion. We thank Dr. Gottfried Otting of the Australian National University for providing the first batch of C3 label, and the Vanderbilt synthesis core for subsequently reproducing the labels in-house. The cysless T4L construct was a kind gift from the Mchaourab lab at Vanderbilt. This research used resources of the Advanced Photon Source, a U.S. Department of Energy (DOE) Office of Science User Facility operated for the DOE Office of Science by Argonne National Laboratory under Contract No. DE-AC02-06CH11357. Use of the LS-CAT Sector 21 was supported by the Michigan Economic Development Corporation and the Michigan Technology Tri-Corridor (Grant 085P1000817).

## Author contributions

E.O. and J.M. conceived the idea. E.O, S.G. and H.D. planned experimental procedure for site-directed lanthanide spin labeling. E.O designed mutants and constructs, transformed, expressed, and purified the proteins, and performed click chemistry reactions. E.O. and K.L. optimized mutant protein preparation and click chemistry conditions. E.O. performed and analyzed results from NMR, X-ray crystallography, and circular dichroism experiments. E.O. analyzed the paramagnetic data. E.O. and J.H. determined the crystal structure presented in this work. E.O. performed the computational calculations and analyzed the data with guidance from G.K., H.S., and J.M. The paper was written by E.O. with input from all authors. This work was supervised by GK and JM. The final manuscript has been approved by all authors.

## Supplementary Figures

**Supplementary Figure 1.**
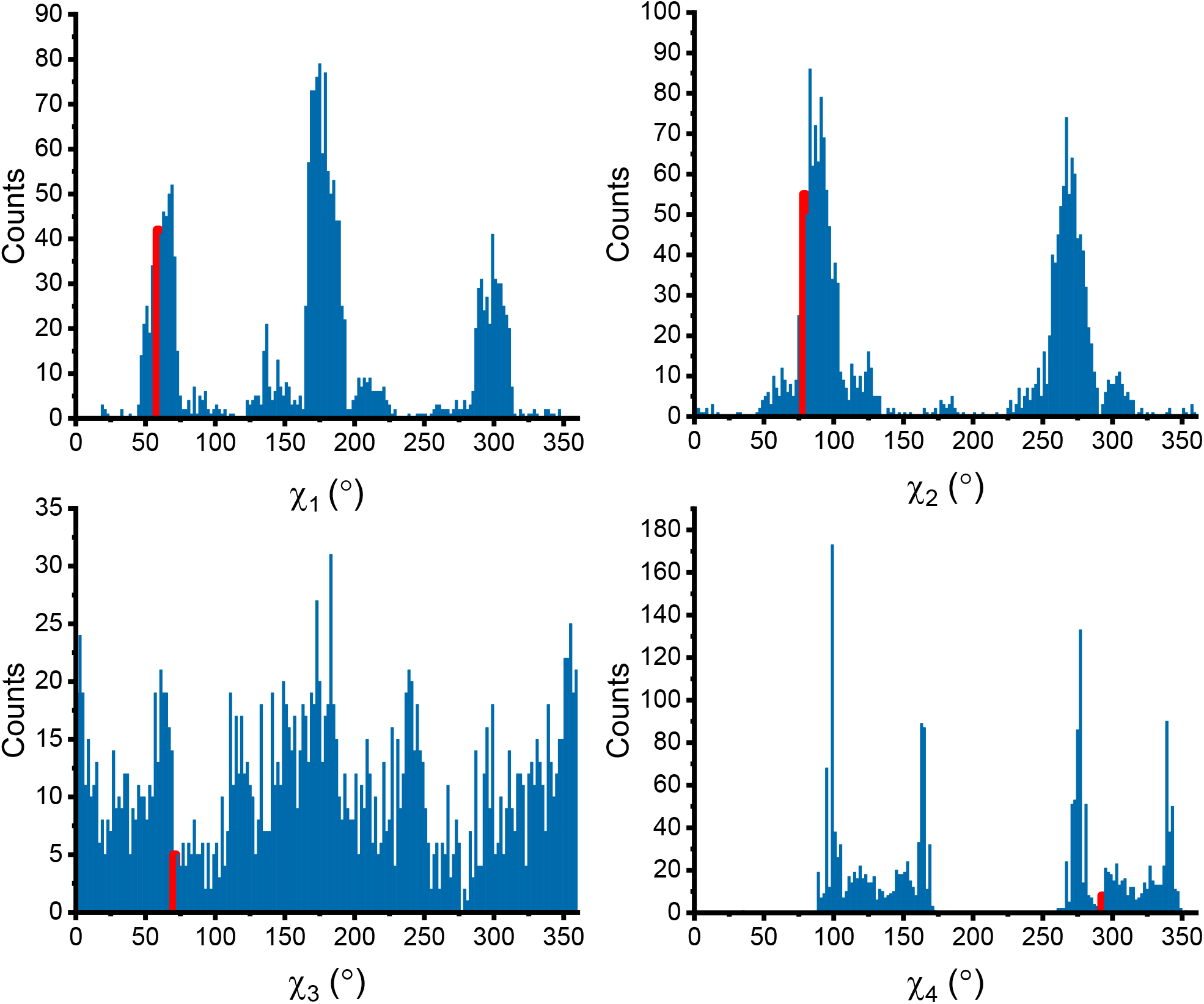
Distribution of rotamers of the pAzF-C3 spin label. The distribution of rotamers for χ1 to χ4 of the pAzF-C3 spin label computed with the BCL conformer generator. The chi angles observed in the crystal structure are indicated on each plot with a red line.

**Supplementary Figure 2.**
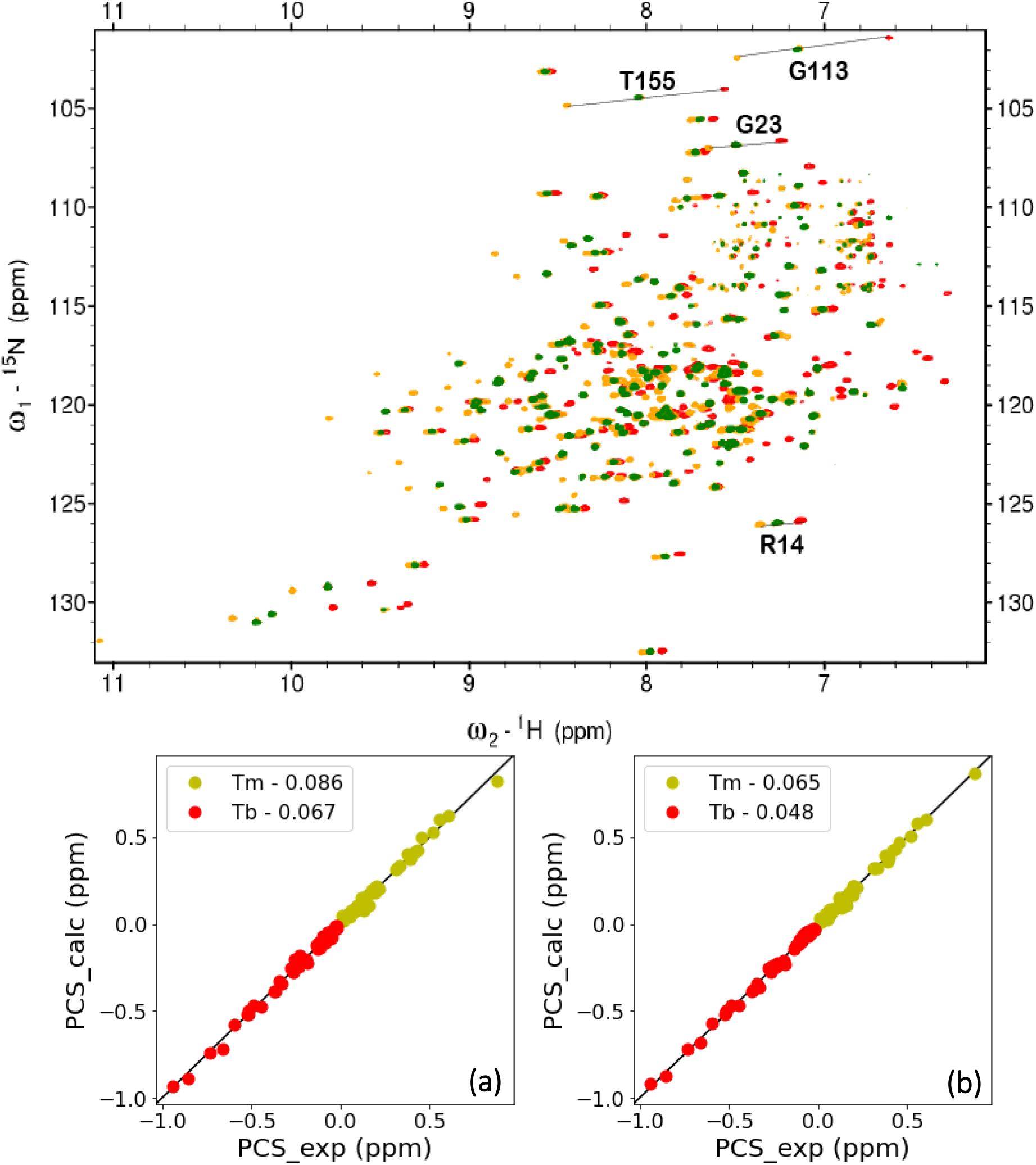
PCS dataset at T4L position 119. Top panel: Superimposition of 2D [^1^H-^15^N]-IPAP spectra of T4L-pAzF119 in complex with one diamagnetic C3-Y^3+^ (green), or paramagnetic C3-Tm^3+^ (yellow), and C3-Tb^3+^ (red). Lines indicate a selection of observed PCSs. PCSs were too small to measure in T4L-pAzF119-C3-Yb^3+^ spectrum (not shown). Bottom panel: The PCS data for both backbone amide protons and nitrogens obtained with Tm^3+^ (yellow), and Tb^3+^ (red) coordinated to the C3 label were fitted globally to (a) the closed structure of T4L (PDB ID: 2LZM) and (b) the open structure of T4L (PDB ID: 172L). The PCS Q-factors between the observed and calculated values are given.

**Supplementary Figure 3.**
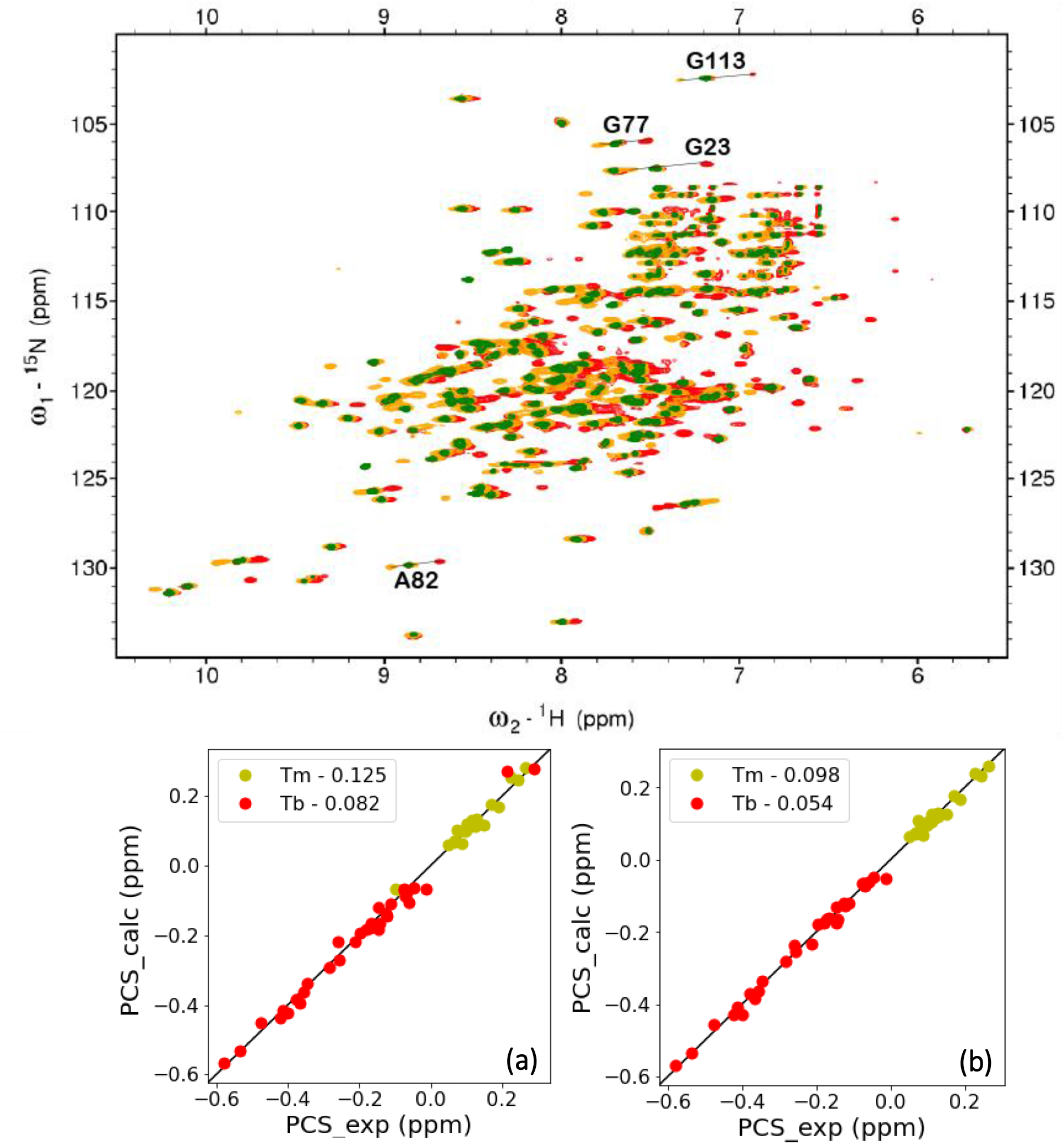
PCS dataset at T4L position 131. Top panel: Superimposition of 2D [^1^H-^15^N]-IPAP spectra of T4L-pAzF131 in complex with one diamagnetic C3-Y^3+^ (green), or paramagnetic C3-Tm^3+^ (yellow), and C3-Tb^3+^ (red). Lines indicate a selection of observed PCSs. PCSs were too small to measure in T4L-pAzF119-C3-Yb^3+^ spectrum (not shown). Bottom panel: The PCS data for both backbone amide protons and nitrogens obtained with Tm^3+^ (yellow), and Tb^3+^ (red) coordinated to the C3 label were fitted globally to (a) the closed structure of T4L (PDB ID: 2LZM) and (b) the open structure of T4L (PDB ID: 172L). The PCS Q-factors between the observed and calculated values are given.

**Supplementary Figure 4.**
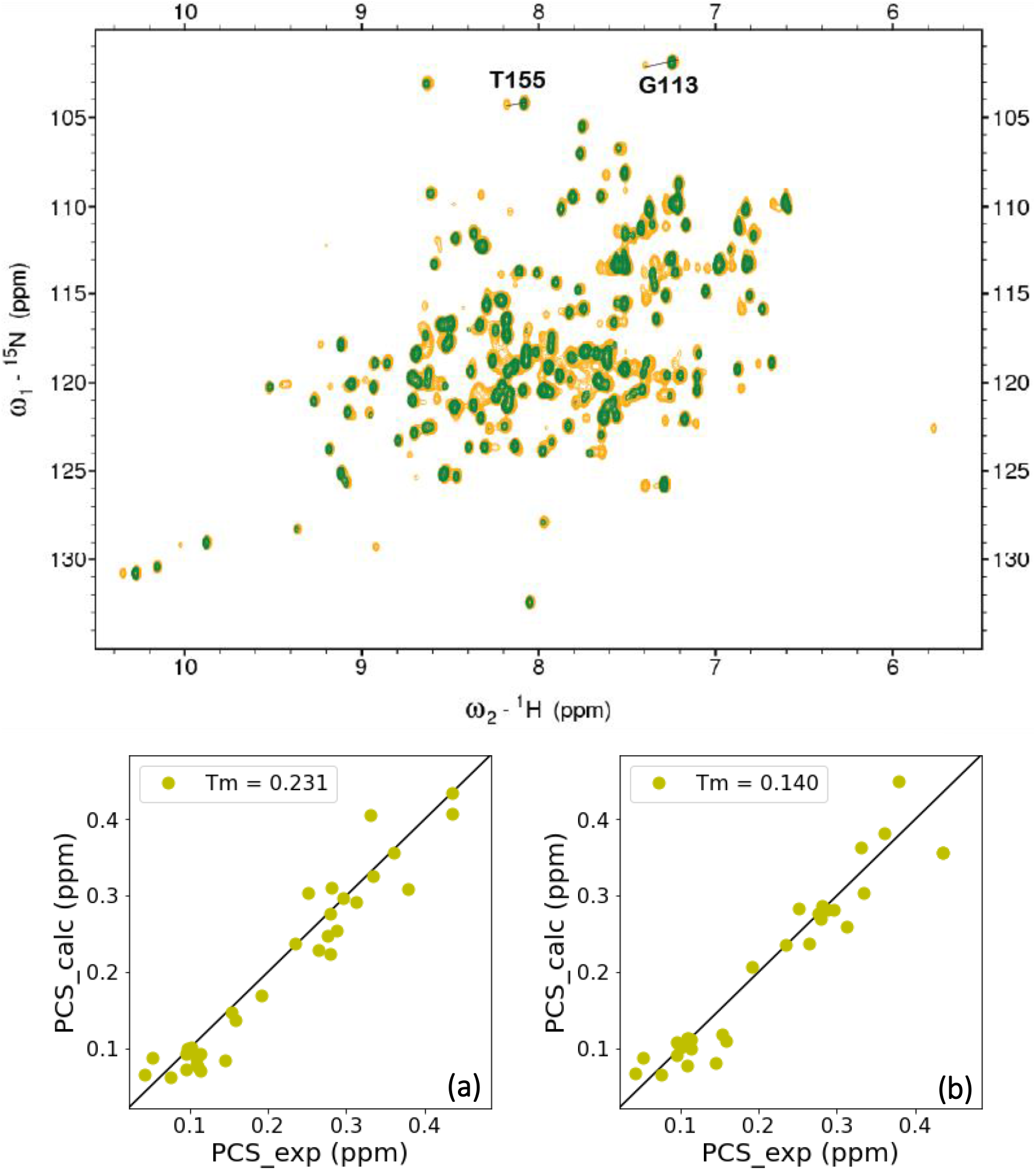
PCS dataset at T4L position 44. Top panel: Superimposition of 2D [^1^H-^15^N]-HSQC spectra of T4L-pAzF44 in complex with one diamagnetic C3-Y^3+^ (green), or paramagnetic C3-Tm^3+^ (yellow). Lines indicate a selection of observed PCSs. PCSs were too small to measure in T4L-pAzF119-C3-Yb^3+^ spectrum (not shown), and most peaks were wiped out in T4L-pAzF-C3-Tb^3+^ (not shown). Bottom panel: The PCS data for both backbone amide protons and nitrogens obtained with Tm^3+^ (yellow) coordinated to the C3 label were fitted globally to (a) the closed structure of T4L (PDB ID: 2LZM) and (b) the open structure of T4L (PDB ID: 172L). The PCS Q-factors between the observed and calculated values are given.

**Supplementary Figure 5.**
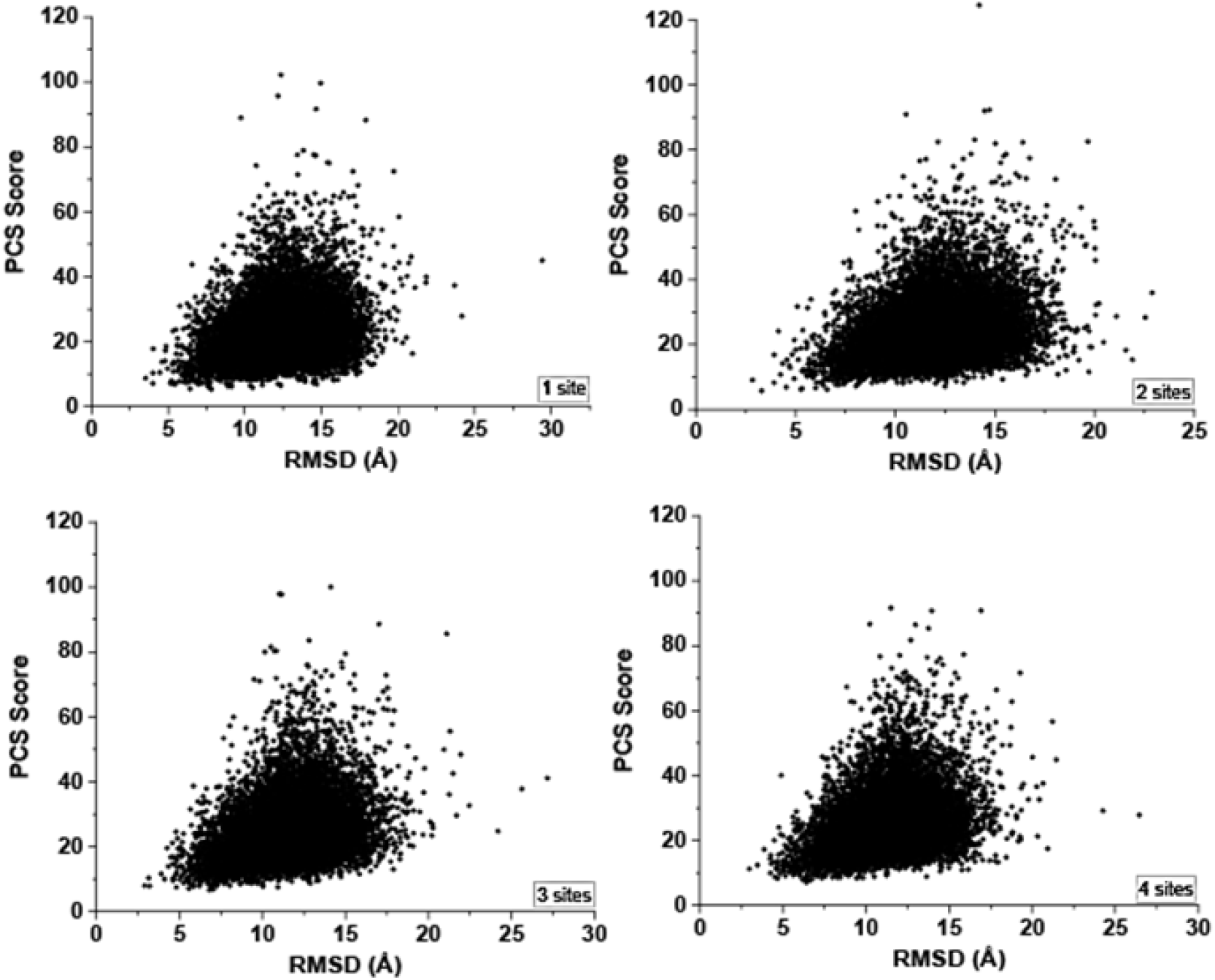
PCS score-vs-RMSD plots for models generated and rescored with PCS data from one, two, three, or four labeling sites. Final model selection was conducted after rescoring decoys with PCSs. Models generated with PCSs at multiple sites resulted in sampling of models closer to the native. Runs including data from four PCS sites yielded the lowest RMSD models from the native, while runs with no PCS data yielded models with the highest RMSD from the native.

**Supplementary Figure 6.**
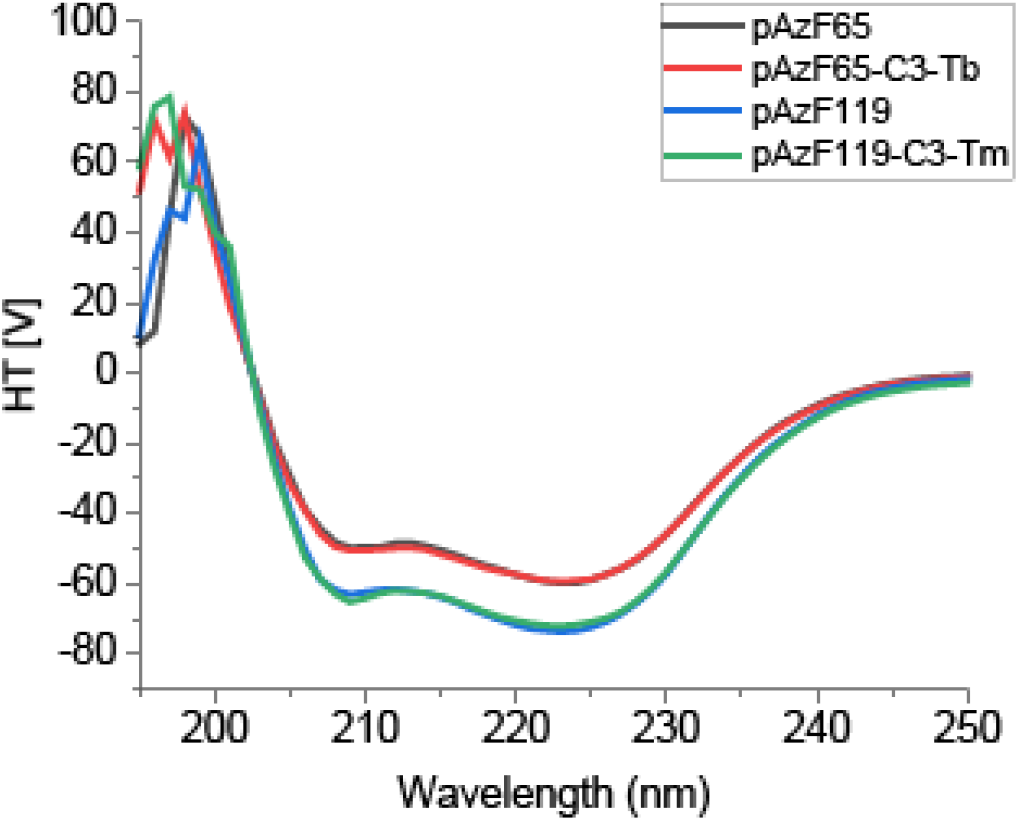
Circular Dichroism spectra show C3-labeled T4L mutants maintain α-helical nature. CD spectra are shown for selected mutants. T4L pAzF mutant before lanthanide labeling was used a reference (black line for T4LpAzF65, and blue line for T4LpAzF119). All lanthanide-labeled mutants maintained α-helical profile expected for T4L, as shown here for T4LpAzF65 labeled with C3-Tb^3+^ (pAzF65-C3-Tb, red) and T4LpAzF119 labeled with C3-Tm^3+^ (pAzF119-C3-Tm, green).

**Supplementary Figure 7.**
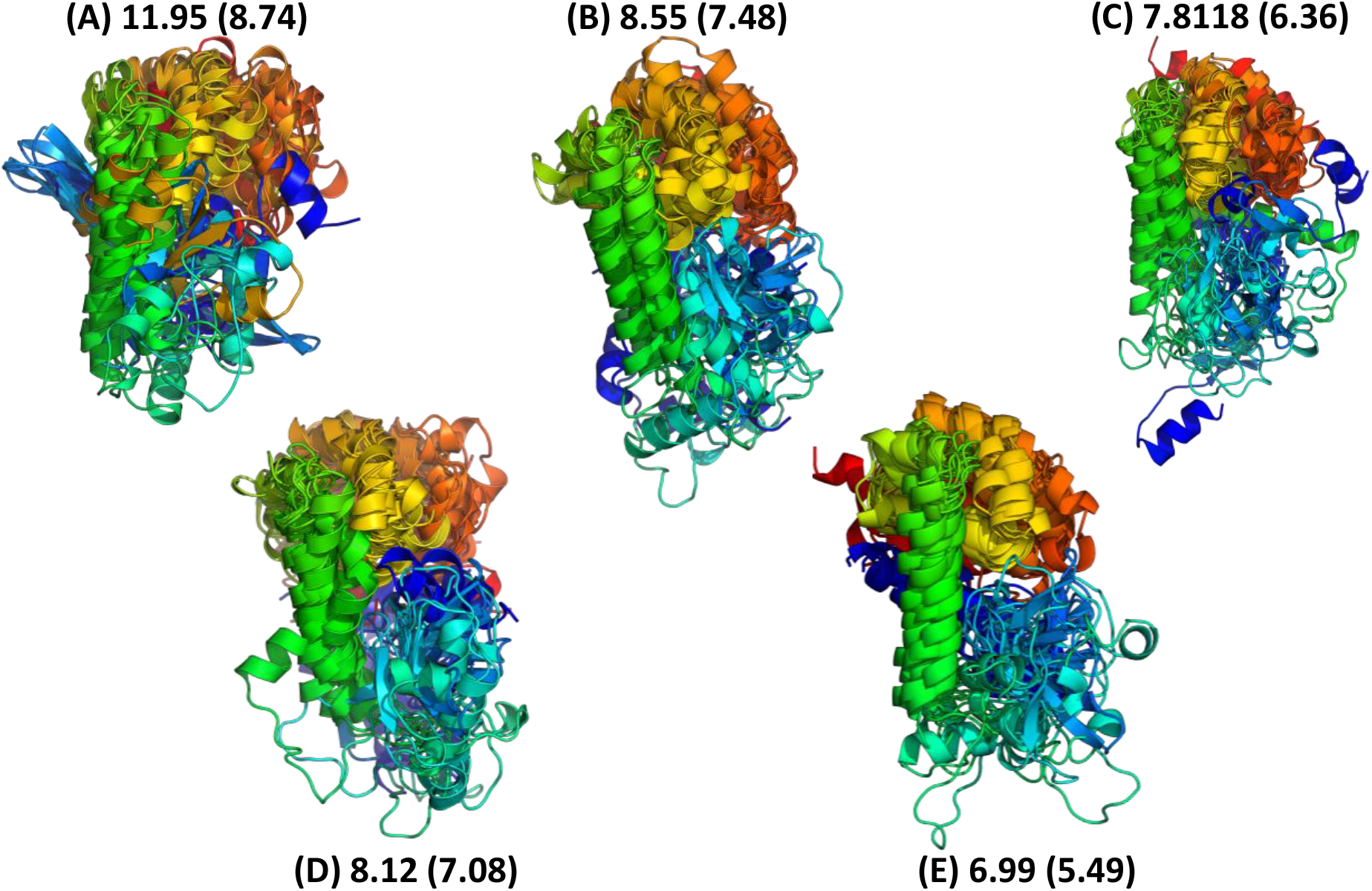
Top models selected after PCS rescoring show models approaching convergence for lowest scoring models. After rescoring models with PCSs, the top 10 models with lowest combined ROSETTA + PCS score showed better convergence when PCSs were used. Improvement was also seen in sampling the more flexible domain when PCSs at multiple sites were included. The average of all pairwise RMSDs in the ensemble is given for the full-length protein, and also for the core α-helical regions in parentheses. Ensembles show models generated with no PCSs (A), PCSs from 1 site (B), 2 sites (C), 3 sites (D), and 4 sites (E).

**Supplementary Table 1:** PCSs (in ppm) measured at site 65 (with C3-Yb^3+^)

|  |  |  |
| --- | --- | --- |
| 14 | H | 0.267 |
| 15 | H | 0.227 |
| 16 | H | 0.251 |
| 18 | H | 0.249 |
| 23 | H | 0.081 |
| 26 | H | 0.247 |
| 27 | H | 0.421 |
| 28 | H | 0.402 |
| 29 | H | 0.430 |
| 31 | H | 0.386 |
| 32 | H | 0.285 |
| 33 | H | 0.338 |
| 39 | H | 0.092 |
| 41 | H | 0.11 |
| 42 | H | 0.182 |
| 55 | H | 0.186 |
| 56 | H | 0.291 |
| 58 | H | 0.357 |
| 60 | H | 0.254 |
| 63 | H | 0.605 |
| 71 | H | 0.113 |
| 80 | H | -0.183 |
| 81 | H | -0.133 |
| 82 | H | -0.087 |
| 85 | H | -0.089 |
| 87 | H | -0.006 |
| 89 | H | -0.073 |
| 91 | H | -0.055 |
| 92 | H | -0.033 |
| 96 | H | -0.036 |
| 107 | H | -0.015 |
| 108 | H | -0.076 |
| 109 | H | -0.064 |
| 113 | H | -0.033 |
| 138 | HE1 | 0.014 |
| 142 | H | 0.016 |
| 164 | H | 0.033 |
| 14 | N | 0.217 |
| 15 | N | 0.272 |
| 16 | N | 0.23 |
| 18 | N | 0.254 |
| 23 | N | 0.066 |
| 26 | N | 0.274 |
| 27 | N | 0.39 |
| 28 | N | 0.441 |
| 29 | N | 0.416 |
| 31 | N | 0.354 |
| 32 | N | 0.326 |
| 33 | N | 0.456 |
| 39 | N | 0.112 |
| 41 | N | 0.141 |
| 42 | N | 0.162 |
| 55 | N | 0.197 |
| 56 | N | 0.28 |
| 58 | N | 0.415 |
| 60 | N | 0.298 |
| 63 | N | 0.72 |
| 71 | N | 0.012 |
| 80 | N | -0.173 |
| 81 | N | -0.14 |
| 82 | N | -0.09 |
| 85 | N | -0.12 |
| 87 | N | -0.063 |
| 89 | N | -0.069 |
| 91 | N | 0.026 |
| 92 | N | -0.044 |
| 96 | N | -0.038 |
| 107 | N | -0.021 |
| 108 | N | -0.109 |
| 109 | N | -0.075 |
| 113 | N | -0.051 |
| 138 | NE1 | 0.003 |
| 142 | N | -0.009 |
| 164 | N | 0.04 |

**Supplementary Table 2:** PCSs (in ppm) measured at site 65 (with C3-Tm^3+^)

|  |  |  |
| --- | --- | --- |
| 12 | H | 0.259 |
| 14 | H | 0.569 |
| 16 | H | 0.577 |
| 18 | H | 0.624 |
| 23 | H | 0.22 |
| 26 | H | 0.630 |
| 27 | H | 1.06 |
| 29 | H | 0.916 |
| 31 | H | 0.942 |
| 33 | H | 0.928 |
| 39 | H | 0.266 |
| 41 | H | 0.32 |
| 42 | H | 0.519 |
| 55 | H | 0.559 |
| 56 | H | 0.789 |
| 58 | H | 0.809 |
| 59 | H | 0.857 |
| 60 | H | 0.432 |
| 63 | H | 1.092 |
| 77 | H | -0.494 |
| 80 | H | -0.355 |
| 81 | H | -0.259 |
| 82 | H | -0.174 |
| 87 | H | -0.119 |
| 89 | H | -0.146 |
| 91 | H | -0.108 |
| 92 | H | -0.086 |
| 108 | H | -0.114 |
| 109 | H | -0.105 |
| 113 | H | -0.052 |
| 142 | H | 0.05 |
| 12 | N | 0.263 |
| 14 | N | 0.498 |
| 16 | N | 0.607 |
| 18 | N | 0.609 |
| 23 | N | 0.183 |
| 26 | N | 0.747 |
| 27 | N | 0.995 |
| 29 | N | 0.895 |
| 31 | N | 0.892 |
| 33 | N | 1.156 |
| 39 | N | 0.304 |
| 41 | N | 0.385 |
| 42 | N | 0.619 |
| 55 | N | 0.588 |
| 56 | N | 0.753 |
| 58 | N | 0.925 |
| 59 | N | 0.756 |
| 60 | N | 0.529 |
| 63 | N | 1.308 |
| 77 | N | -0.497 |
| 80 | N | -0.333 |
| 81 | N | -0.28 |
| 82 | N | -0.177 |
| 87 | N | -0.131 |
| 89 | N | -0.013 |
| 91 | N | -0.081 |
| 92 | N | -0.076 |
| 108 | N | -0.167 |
| 109 | N | -0.13 |
| 113 | N | -0.1 |
| 142 | N | 0.014 |

**Supplementary Table 3:** PCSs (in ppm) measured at site 65 (with C3-Tb^3+^)

|  |  |  |
| --- | --- | --- |
| 12 | H | -0.373 |
| 14 | H | -1.03 |
| 16 | H | -1.143 |
| 23 | H | -0.452 |
| 39 | H | -0.585 |
| 41 | H | -0.722 |
| 56 | H | -1.543 |
| 77 | H | 1.541 |
| 80 | H | 0.422 |
| 107 | H | 0.28 |
| 108 | H | 0.398 |
| 155 | H | 0.108 |
| 12 | N | -0.351 |
| 14 | N | -0.983 |
| 16 | N | -1.126 |
| 23 | N | -0.374 |
| 39 | N | -0.677 |
| 41 | N | -0.877 |
| 56 | N | -1.481 |
| 77 | N | 1.448 |
| 80 | N | 0.227 |
| 108 | N | 0.477 |
| 109 | N | 0.35 |
| 155 | N | 0.107 |

**Supplementary Table 4:** PCSs (in ppm) measured at site 119 (with C3-Tm^3+^)

|  |  |  |
| --- | --- | --- |
| 11 | H | 0.157 |
| 12 | H | 0.095 |
| 14 | H | 0.055 |
| 20 | H | 0.034 |
| 21 | H | 0.051 |
| 23 | H | 0.030 |
| 30 | H | 0.069 |
| 32 | H | 0.044 |
| 71 | H | 0.086 |
| 74 | H | 0.081 |
| 77 | H | 0.049 |
| 89 | H | 0.457 |
| 104 | H | 0.177 |
| 106 | H | 0.174 |
| 107 | H | 0.136 |
| 110 | H | 0.124 |
| 113 | H | 0.334 |
| 114 | H | 0.556 |
| 138 | HE1 | 0.219 |
| 142 | H | 0.152 |
| 155 | H | 0.399 |
| 156 | H | 0.431 |
| 157 | H | 0.316 |
| 158 | HE1 | 0.197 |
| 164 | H | 0.098 |
| 11 | N | 0.143 |
| 12 | N | 0.128 |
| 14 | N | 0.038 |
| 20 | N | 0.007 |
| 21 | N | 0.057 |
| 23 | N | 0.031 |
| 30 | N | 0.075 |
| 32 | N | 0.028 |
| 71 | N | 0.066 |
| 74 | N | 0.061 |
| 77 | N | 0.019 |
| 89 | N | 0.518 |
| 104 | N | 0.196 |
| 106 | N | 0.154 |
| 107 | N | 0.122 |
| 110 | N | 0.118 |
| 113 | N | 0.392 |
| 114 | N | 0.606 |
| 138 | NE1 | 0.201 |
| 142 | N | 0.120 |
| 155 | N | 0.378 |
| 156 | N | 0.418 |
| 157 | N | 0.312 |
| 158 | NE1 | 0.193 |
| 164 | N | 0.087 |

**Supplementary Table 5:** PCSs (in ppm) measured at site 119 (with C3-Tb^3+^)

|  |  |  |
| --- | --- | --- |
| 12 | H | -0.138 |
| 14 | H | -0.089 |
| 20 | H | -0.058 |
| 21 | H | -0.083 |
| 23 | H | -0.055 |
| 29 | H | -0.090 |
| 30 | H | -0.104 |
| 32 | H | -0.071 |
| 33 | H | -0.047 |
| 51 | H | -0.019 |
| 56 | H | -0.023 |
| 59 | H | -0.045 |
| 60 | H | -0.059 |
| 71 | H | -0.121 |
| 74 | H | -0.119 |
| 77 | H | -0.071 |
| 89 | H | -0.658 |
| 104 | H | -0.265 |
| 106 | H | -0.266 |
| 107 | H | -0.215 |
| 110 | H | -0.194 |
| 113 | H | -0.517 |
| 126 | HE1 | -0.850 |
| 138 | HE1 | -0.346 |
| 142 | H | -0.256 |
| 155 | H | -0.485 |
| 156 | H | -0.521 |
| 157 | H | -0.372 |
| 164 | H | -0.129 |
| 14 | N | -0.098 |
| 20 | N | -0.053 |
| 23 | N | -0.046 |
| 29 | N | -0.098 |
| 30 | N | -0.096 |
| 51 | N | -0.027 |
| 60 | N | -0.073 |
| 71 | N | -0.128 |
| 74 | N | -0.108 |
| 77 | N | -0.027 |
| 89 | N | -0.733 |
| 104 | N | -0.278 |
| 106 | N | -0.234 |
| 107 | N | -0.201 |
| 110 | N | -0.183 |
| 113 | N | -0.592 |
| 126 | NE1 | -0.942 |
| 138 | NE1 | -0.331 |
| 142 | N | -0.225 |
| 155 | N | -0.446 |
| 156 | N | -0.519 |
| 157 | N | -0.363 |
| 164 | N | -0.114 |

**Supplementary Table 6:** PCSs (in ppm) measured at site 131 (with C3-Tm^3+^)

|  |  |  |
| --- | --- | --- |
| 71 | H | 0.068 |
| 77 | H | 0.097 |
| 79 | H | 0.127 |
| 80 | H | 0.112 |
| 82 | H | 0.101 |
| 87 | H | 0.265 |
| 89 | H | 0.245 |
| 106 | H | 0.126 |
| 113 | H | 0.150 |
| 138 | HE1 | 0.189 |
| 158 | HE1 | 0.089 |
| 164 | H | 0.099 |
| 71 | N | 0.066 |
| 77 | N | 0.082 |
| 79 | N | 0.127 |
| 80 | N | 0.109 |
| 82 | N | 0.114 |
| 89 | N | 0.226 |
| 106 | N | 0.075 |
| 113 | N | 0.111 |
| 138 | NE1 | 0.170 |
| 158 | NE1 | 0.050 |
| 164 | N | -0.084 |

**Supplementary Table 7:**
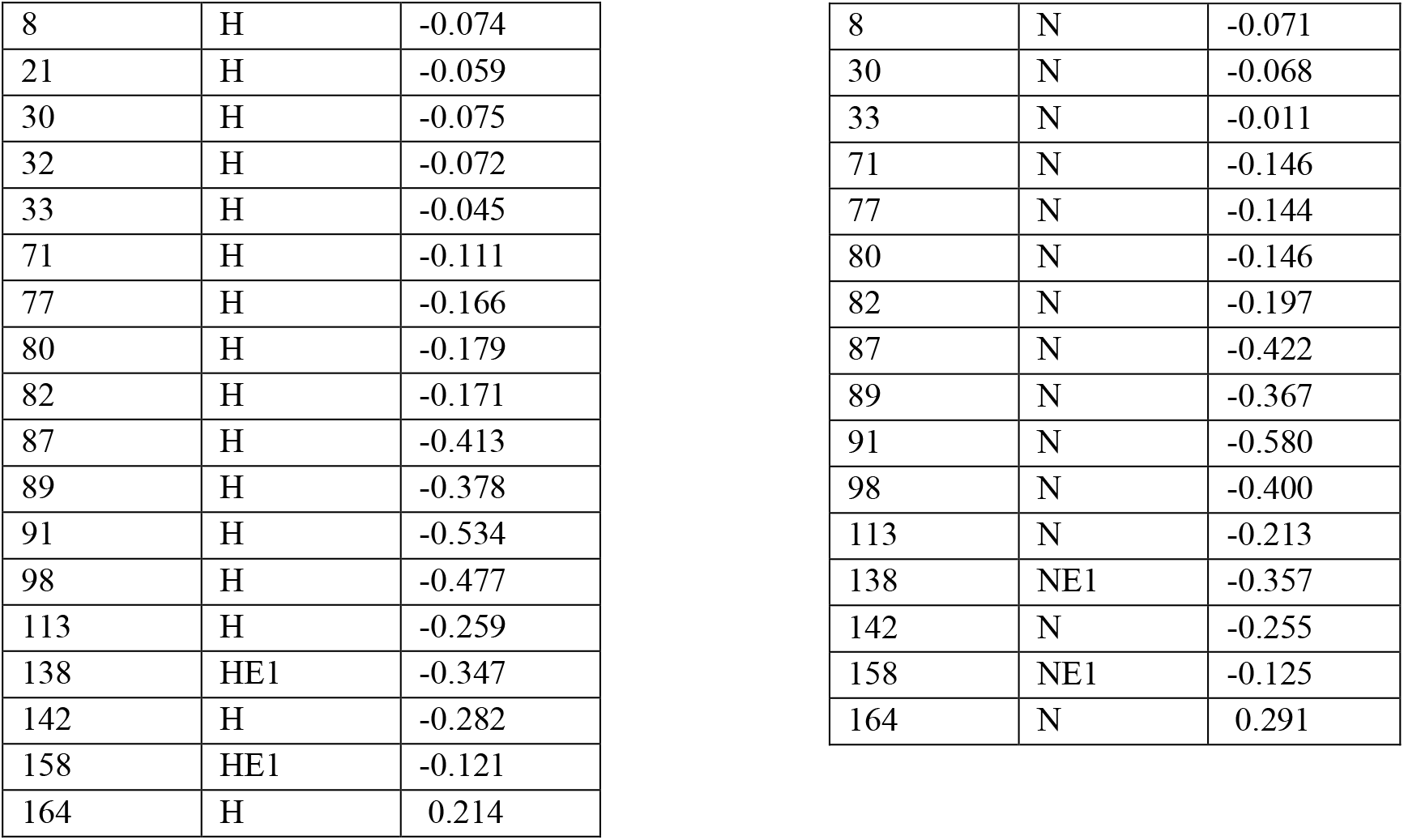
PCSs (in ppm) measured at site 131 (with C3-Tb^3+^)

|  |  |  |
| --- | --- | --- |
| 8 | H | -0.074 |
| 21 | H | -0.059 |
| 30 | H | -0.075 |
| 32 | H | -0.072 |
| 33 | H | -0.045 |
| 71 | H | -0.111 |
| 77 | H | -0.166 |
| 80 | H | -0.179 |
| 82 | H | -0.171 |
| 87 | H | -0.413 |
| 89 | H | -0.378 |
| 91 | H | -0.534 |
| 98 | H | -0.477 |
| 113 | H | -0.259 |
| 138 | HE1 | -0.347 |
| 142 | H | -0.282 |
| 158 | HE1 | -0.121 |
| 164 | H | 0.214 |
| 8 | N | -0.071 |
| 30 | N | -0.068 |
| 33 | N | -0.011 |
| 71 | N | -0.146 |
| 77 | N | -0.144 |
| 80 | N | -0.146 |
| 82 | N | -0.197 |
| 87 | N | -0.422 |
| 89 | N | -0.367 |
| 91 | N | -0.580 |
| 98 | N | -0.400 |
| 113 | N | -0.213 |
| 138 | NE1 | -0.357 |
| 142 | N | -0.255 |
| 158 | NE1 | -0.125 |
| 164 | N | 0.291 |

**Supplementary Table 8:** PCSs (in ppm) measured at site 44 (with C3-Tm^3+^)

|  |  |  |
| --- | --- | --- |
| 8 | H | 0.331 |
| 20 | H | 0.379 |
| 21 | H | 0.287 |
| 23 | H | 0.264 |
| 39 | H | 0.276 |
| 59 | H | 0.463 |
| 82 | H | 0.108 |
| 89 | H | 0.114 |
| 91 | H | 0.11 |
| 92 | H | 0.109 |
| 97 | H | 0.192 |
| 106 | H | 0.296 |
| 107 | H | 0.334 |
| 113 | H | 0.153 |
| 126 | H | 0.076 |
| 155 | H | 0.096 |
| 156 | H | 0.101 |
| 157 | H | 0.103 |
| 8 | N | 0.435 |
| 20 | N | 0.313 |
| 23 | N | 0.279 |
| 39 | N | 0.28 |
| 59 | N | 0.435 |
| 77 | N | 0.234 |
| 82 | N | 0.145 |
| 91 | N | 0.095 |
| 104 | N | 0.36 |
| 106 | N | 0.252 |
| 107 | N | 0.281 |
| 113 | N | 0.158 |
| 126 | N | 0.043 |
| 155 | N | 0.052 |
| 156 | N | 0.114 |
| 157 | N | 0.098 |

**Supplementary Table 9:** PCS Q-factors calculated for N-terminal region in T4L.

| Position | 65 | 131 | 119 | 44 |
| --- | --- | --- | --- | --- |
| C3-Yb<sup>3+</sup> | 0.067 (0.076)<sup>i</sup> | N/A | N/A | N/A |
| C3-Tm<sup>3+</sup> | 0.045 (0.052) | 0.086 (0.122) | 0.16 (0.172) | 0.082 (0.984) |
| C3-Tb<sup>3+</sup> | 0.039 (0.054) | No PCSs<sup>ii</sup> | 0.087 (0.174) | N/A |
| Average | 0.050 (0.061) | 0.086 (0.122) | 0.124 (0.173) | 0.082 (0.984) |

<sup>i</sup> PCS Q-factors reported for open form of T4L, and in parenthesis are for closed form of T4L
<sup>ii</sup> There were not enough PCSs obtained from the N-terminal region with C3-Tb<sup>3+</sup> at position 131 to fit independently.

